# Flexible resonance in prefrontal networks with strong feedback inhibition

**DOI:** 10.1101/364729

**Authors:** Jason S. Sherfey, Salva Ardid, Joachim Hass, Michael E. Hasselmo, Nancy J. Kopell

## Abstract

Oscillations are ubiquitous features of brain dynamics that undergo task-related changes in synchrony, power, and frequency. The impact of those changes on target networks is poorly understood. In this work, we used a biophysically detailed model of prefrontal cortex (PFC) to explore the effects of varying the spike rate, synchrony, and waveform of strong oscillatory inputs on the behavior of cortical networks driven by them. Interacting populations of excitatory and inhibitory neurons with strong feedback inhibition are inhibition-based network oscillators that exhibit resonance (i.e., larger responses to preferred input frequencies). We quantified network responses in terms of mean firing rates and the population frequency of network oscillation; and characterized their behavior in terms of the natural response to asynchronous input and the resonant response to oscillatory inputs. We show that strong feedback inhibition causes the PFC to generate internal (natural) oscillations in the beta/gamma frequency range (>15 Hz) and to maximize principal cell spiking in response to external oscillations at slightly higher frequencies. Importantly, we found that the fastest oscillation frequency that can be relayed by the network maximizes local inhibition and is equal to a frequency even higher than that which maximizes the firing rate of excitatory cells; we call this phenomenon population frequency resonance. This form of resonance is shown to determine the optimal driving frequency for suppressing responses to asynchronous activity. Lastly, we demonstrate that the natural and resonant frequencies can be tuned by changes in neuronal excitability, the duration of feedback inhibition, and dynamic properties of the input. Our results predict that PFC networks are tuned for generating and selectively responding to beta- and gamma-rhythmic signals due to the natural and resonant properties of inhibition-based oscillators. They also suggest strategies for optimizing transcranial stimulation and using oscillatory networks in neuromorphic engineering.

**Author Summary:** The prefrontal cortex (PFC) flexibly encodes task-relevant representations and outputs biases to mediate higher cognitive functions. The relevant neural ensembles undergo task-related changes in oscillatory dynamics at beta- and gamma frequencies. Using a computational model of the PFC network, we show that strong feedback inhibition causes the PFC to generate internal oscillations and to prefer external oscillations at similar frequencies. The precise frequencies that are generated and preferred can be flexibly tuned by varying the synchrony and strength of input network activity, the level of background excitation, and neuromodulation of intrinsic ion currents. We also show that the peak output frequency in response to external oscillations, which depends on the synchrony and strength of the input as well as the strong inhibitory feedback, is faster than the internally generated frequency, and that this difference enables exclusive response to oscillatory inputs. These properties enable changes in oscillatory dynamics to govern the selective processing and gating of task-relevant signals in service of cognitive control.

## Introduction

Oscillatory neural activity is a common feature of brain dynamics. In vitro experiments have demonstrated that different brain regions can produce network oscillations at different frequencies [1,2]. In vivo experiments have shown that field potential oscillations in prefrontal cortex (PFC) at beta-(15-35Hz) and gamma-(35-80Hz) frequencies undergo task-related modulations in their power [3] and synchrony [4] and that multiple frequencies can appear in the same region [5,6]. Despite the wealth of experimental evidence suggesting changes in oscillation frequency and synchrony are functionally significant, little remains known about the mechanisms by which they affect processing in downstream networks (but see [7]). In this paper, we will explore the natural, resonant, and competitive dynamics of PFC networks and how the task-modulated properties of oscillatory signals affect those dynamics.

Neural systems at multiple scales are known to exhibit larger responses to oscillatory inputs at preferred (resonant) frequencies. For instance, neurons can exhibit resonance in subthreshold voltage fluctuations [8,9], and networks can exhibit resonance in the amplitude of suprathreshold instantaneous firing rates of self-inhibiting interneurons (INs) [10] and reciprocally connected populations of principal cells (PCs) and INs [11]. Given weak inputs, these systems often exhibit response amplitudes that scale linearly with the input, and they oscillate with the same frequency as the input. In the linear regime, analytical methods can be applied to fully characterize network responses [11]. However, the results of such analyses no longer hold when inputs are strong and responses become strongly nonlinear.

Neural models of fast network oscillations, like those observed in PFC, often involve populations of cells receiving strong feedback inhibition to synchronize the network and strong excitatory input to drive the oscillation [6]. Under a constant (possibly noisy) tonic input, self-inhibiting populations of INs and reciprocally connected PC and IN populations can generate (natural) gamma-frequency network oscillations, termed ING [1,12] and PING [12,13], respectively. Due to the strong input and oscillatory response to a tonic drive, the earlier work on network resonance does not extend to these inhibition-based oscillators.

In this article, we present a numerical study of the natural and resonant behavior of an inhibition-based PC/IN network oscillator. In contrast to the linear regime, we will show that the frequency of the network oscillation equals the input up to a maximal frequency, above which, it decreases; we call this phenomenon population frequency resonance. We will show that different input frequencies maximize inhibition and excitation when inputs are strong and that the population frequency peaks when inhibition is maximized. The importance of this phenomenon will be demonstrated by showing that the optimal driving frequency for suppressing responses to asynchronous input is that which maximizes population frequency and not that which maximizes the firing rate of excitatory PCs. Finally, we will show how network resonance depends on dynamic, task-modulated properties of the input as well as intrinsic properties of the resonant network. Our quantitative results identify mechanisms that are not model specific, as will be shown by analogous simulations in a Hodgkin-Huxley type model of PFC and a generic integrate-and-fire network model that exhibit qualitatively similar behavior.

The paper will begin with a characterization of network responses to asynchronous and oscillatory inputs. Responses will be characterized in terms of firing rate and population frequency, and then the latter will be shown to determine the maximal suppression of asynchronous activity. The dependence of response on experimentally-motivated input parameters will be described. Finally, the paper will end with a discussion of the functional relevance of these findings for flexible neural processing.

## Results

We explored the impact of modulating task-related signals on cortical processing using an experimentally-constrained, Hodgkin-Huxley type network model of prefrontal cortex (PFC). Principal cell (PC) activity in PFC can be interpreted as a bias signal that mediates higher cognitive functions. The model represents a deep output layer of reciprocally-connected PCs and fast spiking interneurons (INs) that provide strong feedback inhibition. The network was driven by collections of independent spike trains modeling upstream activity in populations of excitatory cells. The input spike trains were either asynchronous with constant rate or mediated by an oscillatory modulation of the rate. Rhythmic activity is considered task-relevant [14] while the asynchronous activity is task-irrelevant (see Discussion for further considerations). We studied how the network behavior varies with task-related changes in synchrony, frequency, and strength of periodic inputs, and how that behavior relates to the response driven by equal-strength, asynchronous activity.

### Response of the PC/IN network with spiking input

#### PC/IN networks generate non-sinusoidal rhythms in response to asynchronous spiking

Disconnected PCs respond to a tonic input of asynchronous spiking (Fig 1A) with asynchronous responses (Fig 1Bi). Similar to PING oscillations driven by a noisy tonic drive [13], PC/IN networks with increasingly strong feedback inhibition respond to asynchronous spiking with an increasingly periodic modulation of instantaneous firing rate (iFR); the input strength-dependent frequency of the response to an asynchronous input will be called the natural frequency of the network, *f_N_*, for a given input strength (Figs 1Bii-iii; see Table 1 for definitions of the symbols used throughout this paper). Notably, the response of the PC/IN network is pulsatile (Fig 1Biii, iFR trace) because of how quickly INs silence the PC population and the longer time required for PC-synchronizing inhibition to decay; this results in spike trains that are more synchronous than would occur in an oscillation with sinusoidal rate-modulation; this form of spike synchrony in the input will be shown below to have significant consequences for responses in downstream networks. As with all inhibition-based oscillators, the rhythm period increases with the duration of inhibition ( [6], S1 Fig), and the network remains silent within a cycle as long as the inhibition remains sufficiently strong. Over time, this results in the PC/IN network outputting periodic volleys of spikes (i.e., pulse packets) separated by periods of inhibition.

**Figure 1.**
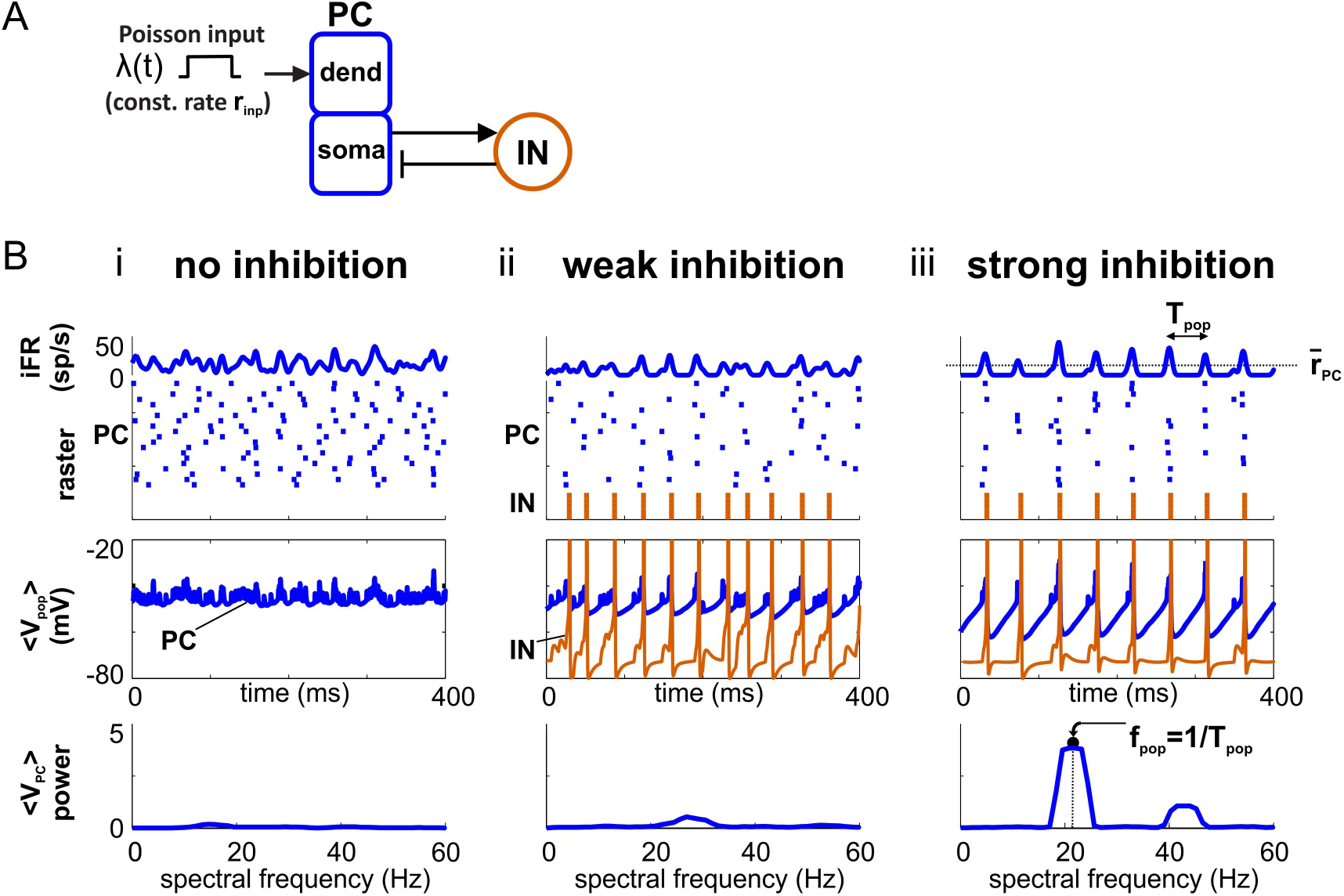
Strong feedback inhibition produces natural oscillation in PC/IN network. (**A**) Diagram showing feedforward excitation from external (independent) Poisson spike inputs to 20 excitatory principal cells (PCs) receiving feedback inhibition from 5 inhibitory interneurons (INs). See Fig 7A for details. (**B**) Simulations showing the network switching from an asynchronous to oscillatory state with natural oscillation as the strength of feedback inhibition is increased.

**Table 1.** Meaning of symbols used in the study of resonance and gating.

| Symbol | Description |
| --- | --- |
| $\lambda(t)$ | Instantaneous input rate of Poisson process (kHz) |
| $r_{inp}$ | Time-averaged Poisson input rate, $\langle \lambda \rangle_t$ (kHz) |
| $f_{inp}$ | Frequency of Poisson input rate-modulation (Hz) |
| $\delta_{inp}$ | Pulse width of Poisson input with square wave rate-modulation (ms) |
| $iFR$ | Instantaneous output firing rate averaged over principal cells (sp/s) |
| $\bar{r}_{PC}$ | Time-averaged population firing rate of principal cells (sp/s) |
| $\bar{r}_{IN}$ | Time-averaged population firing rate of interneurons (sp/s) |
| $f_{pop}$ | Frequency of output population rhythm (Hz); identical for PCs and INs |
| $f_N$ | Natural frequency (Hz) (i.e., $f_{pop}$ elicited by asynchronous input) |
| $f_R^{PC}$ | $\bar{r}_{PC}$-resonant frequency (Hz) (i.e., input $f_{inp}$ maximizing output $\bar{r}_{PC}$) |
| $f_R^{IN}$ | $\bar{r}_{IN}$-resonant frequency (Hz) (i.e., input $f_{inp}$ maximizing output $\bar{r}_{IN}$) |
| $f_R^{pop}$ | $f_{pop}$-resonant frequency (Hz) (i.e., input $f_{inp}$ maximizing output $f_{pop}$) |

#### Natural and resonant frequencies are different in strongly-driven PC/IN networks

Linear oscillators respond to a sinusoidal input with an amplitude that depends on the input frequency and with a frequency that matches its input frequency. The response of the strongly-driven PC/IN network deviates from this behavior in several respects, and it is at this point that a more careful examination of what is meant by input and output of a PC/IN network is required.

Oscillatory inputs to networks of neurons are often treated as sinusoidal. An important reason for this is that the sinusoidal frequency response of a linear system completely describes the system and contains all the information needed to derive its response to any signal [11]. However, as we have already shown, inhibition-based PC/IN oscillators are non-sinusoidal, and we are investigating a nonlinear (i.e., strong input) regime. Therefore, we will use numerical simulation to investigate the network response to non-sinusoidal inputs, perhaps delivered from other upstream PC/IN oscillators. As an approximation to the kind of periodic pulse packets that PC/IN networks generate, we will explore the effects of periodic Poisson signals with square wave rate-modulation in addition to the more traditional signals with sine wave rate-modulation.

Outputs from neurons and populations of neurons are usually analyzed in terms of firing rates because spikes drive neurotransmission and their rates determine integrated effects on postsynaptic neurons. However, postsynaptic activation of PCs can depend more strongly on the frequency of input population oscillation than the firing rates of presynaptic neurons [10,15,16]. Thus, to characterize outputs in terms of properties that determine downstream effects, we analyzed the collective population frequency (i.e., the frequency of output rate-modulation) in addition to the time- and population-averaged firing rates of PCs and INs (see Methods for more details on the choice of output measures). The population frequency differs from the mean PC firing rate when only a fraction of PCs spike per cycle or PCs spike more than once per cycle on average. If PC/IN networks were like linear oscillators, their frequency would be inherited from the input; however, as we will show in PC/IN networks, the output frequency has its own peak (i.e., exhibits resonance), and its characterization is equally important.

Given this understanding of inputs and outputs for PC/IN networks, we next contrasted the response to an ongoing tonic input of asynchronous spiking with the responses to sinusoidal and high-synchrony, square wave inputs with equal-strength (i.e., equal time-averaged rate *r_inp_*) and frequency *f_inp_* (Fig 2A), in analogy to the tonic and frequency responses for linear oscillators described above. We plotted the mean population firing rates, *r̄_PC_* and *r̄_1N_*, in response to square waves (Fig 2Bi) and sine waves (Fig 2C) as measures of output activity for PC and IN populations, respectively. In the strong input regime, all input frequencies elicited a response. Rhythmic inputs produced greater PC responses than asynchronous inputs for most input frequencies (compare the solid *r̄_PC_* curve to the horizontal dashed line in Figs 2Bi and 2C) because their more synchronous spike trains enabled a larger fraction of more correlated PCs to reach threshold before INs were sufficiently engaged to silence the entire population (Fig 2D-E).

**Figure 2.**
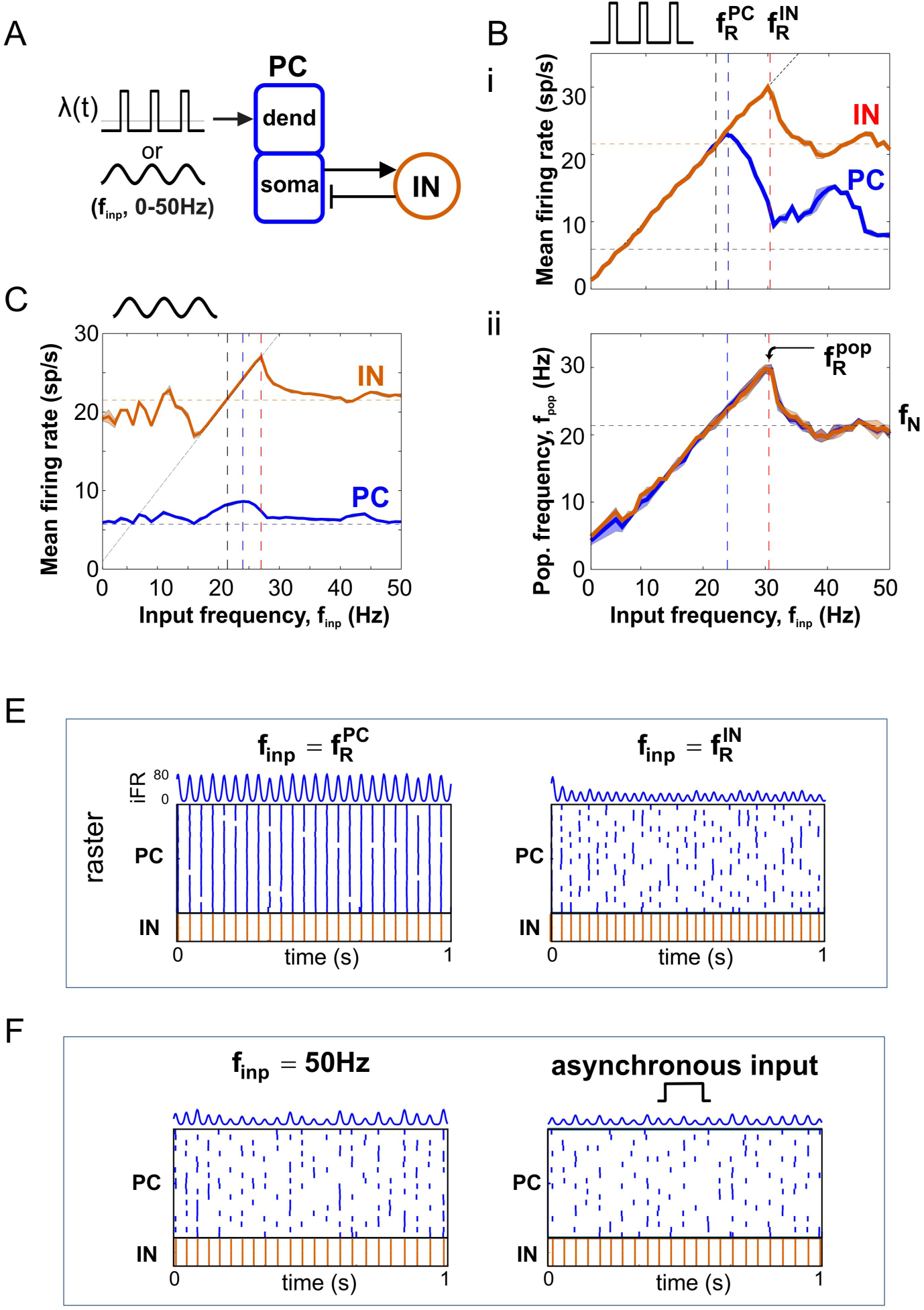
Input frequency-dependent output response profiles. (**A**) Diagram of PFC network receiving sinusoidal or high-synchrony square wave input. (**B**) Response to high-synchrony input. (i) Mean firing rate (FR) profile for PC (blue) and IN (red) populations. Horizontal dashed lines mark the FR response to equal-strength asynchronous input. The diagonal dashed (1:1) line marks where firing rate equals input frequency. (ii) Population frequency profile for PC and IN populations. Peak population frequency occurs at the input frequency maximizing IN activity (i.e., feedback inhibition). Horizontal dashed lines mark the natural frequency in response to asynchronous input. (**C**) FR profile for PC (blue) and IN (red) populations in response to sinusoidal input. Dashed lines mark the same features as in (Bi). (**D**) Spike rasters and PC iFR responses to oscillatory inputs at the PC and IN firing rate resonant frequencies. (**E**) Spike rasters and PC iFR responses to inputs producing network responses paced by internal time constants: (left) asynchronous input, (right) high-frequency input.

Given sinusoidal drive, the fraction of PCs that could spike before being inhibited increased for input frequencies around the natural frequency, *f_N_*, relative to input frequencies far from *f_N_*; the fraction peaked at 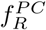 slightly above *f_N_* (Fig 2C, blue curve). Given fixed-mean, high-synchrony square wave drive, all cells spiked on every cycle up to a peak due to the larger instantaneous amplitudes that are present at low frequencies for such inputs (see Methods for a qualitative comparison of *r̄_PC_* response profiles between weak vs. strong and square vs. sine wave inputs). For both input waveforms, *r̄_PC_* peaked at the same 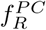 = 24 Hz, and the number of PCs spiking per cycle (i.e., the iFR amplitude) decreased beyond 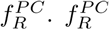 is always ≥ *f_N_* in our model because the correlated spiking of oscillatory inputs produces larger instantaneous drives than the equal-mean asynchronous input while the strength and duration of feedback inhibition on each cycle are the same for both oscillatory and asynchronous inputs. Divergence between natural and resonant frequencies is common in nonlinear [17] and linear systems [8] that show resonance and intrinsic oscillations, except for the harmonic oscillator where they are equivalent. This peak in PC population response to rhythmic inputs with frequencies near *f_N_* depends on matching periodic drives with the rate-limiting time constants of the driven network [18]. The mechanism that determines the precise value of 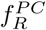 in the inhibition-based PC/IN oscillator is not fully understood (see Discussion). Like 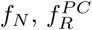 decreases with increasing duration of inhibition (S1 Fig), and the dependence of both on input properties and intrinsic modulatory currents will be presented in later sections.

In contrast to PC firing rates peaking near the natural frequency, *f_N_*, time-averaged IN firing rates continued to increase until input frequency reached 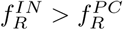 (Fig 2Bi, red curve), where the decreasing number of PCs spiking per cycle became too few to induce IN spiking on each cycle (see next section for more details). Interestingly, this shows that activity of INs driven exclusively by PCs can continue increasing with the frequency of a rhythmic drive to PCs even when PC activity is decreasing, and, consequently, that firing rate resonant frequencies of PC and IN populations can differ. This divergence of 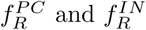 required (1) strong input (S1 Fig), (2) a population of PCs with noisy spiking (S2 Fig), (3) PC → IN synapses that are strong enough for a fraction of PCs to activate INs (S2 Fig), and (4) an IN capable of producing higher firing rates than the PC. For instance, if there is only a single PC (or all PCs spike at the same moment), then the INs can spike only when the PC spikes; thus, the IN spike rate would necessarily decrease with the PC spike rate, and the two would peak for the same input frequency. However, if there is a PC population with noisy spiking and strong PC → IN synapses, then spikes in a subset of the PC population can engage the INs on a given cycle without requiring all cells in the PC population to spike. In this case, even when the time- and population-averaged PC firing rate decreases, INs can continue spiking on every cycle of the input. These qualitative results also hold for leaky integrate-and-fire (LIF) networks (S3 Fig), and the quantitative results hold in a PFC network with 5 times as many PC cells (S4 Fig). Response properties for input frequencies below *f_N_* and above 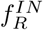 will be shown to depend on input synchrony and strength.

#### Peak oscillation frequency is determined by peak interneuron firing

At the population level, outputs can be further described by the frequency of population oscillation, *f_pop_* (Fig 2Bii); that is, the modulation frequency of the instantaneous population firing rate, or equivalently, the inter-pulse frequency of spike packets emitted by the PC/IN network (see Methods for more details). *f_pop_* profiles also exhibited a peak at a particular input frequency, 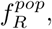 a phenomenon we call population frequency resonance. For square and sine wave inputs to the PFC and LIF networks, the population frequency peaked at 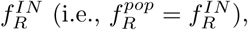 then approached the natural frequency as the response became paced by the network’s internal time constants (i.e., *f_pop_* → *f_N_* as *f_inp_* → ∞) (Fig 2Bii). The population frequency peaked at 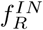 because the PC population could lock to the period of the drive only as long as INs were able to synchronously silence the PC population on each cycle of the input (Fig 2D-E). This yields the possibly counterintuitive result for the inhibition-based PC/IN oscillator that the fastest output oscillation (but not the highest PC firing rate) occurs at the excitatory input frequency that maximizes feedback inhibition.

### Population frequency resonance suppresses response to asynchronous activity

The difference in natural and resonant frequencies in the PC/IN network has at least one functional consequence that we will introduce here. It will serve as motivation for our further exploration of the dependence of natural and resonant frequencies on other properties in the remaining sections below. Consider two parallel pathways driving separate output PC populations that are reciprocally connected to a shared pool of INs (Fig 3A). One pathway (the target) delivers a rhythmic signal to one PC population while the opposing pathway (the distractor) delivers an equal-strength asynchronous signal to the competing PC population.

**Figure 3.**
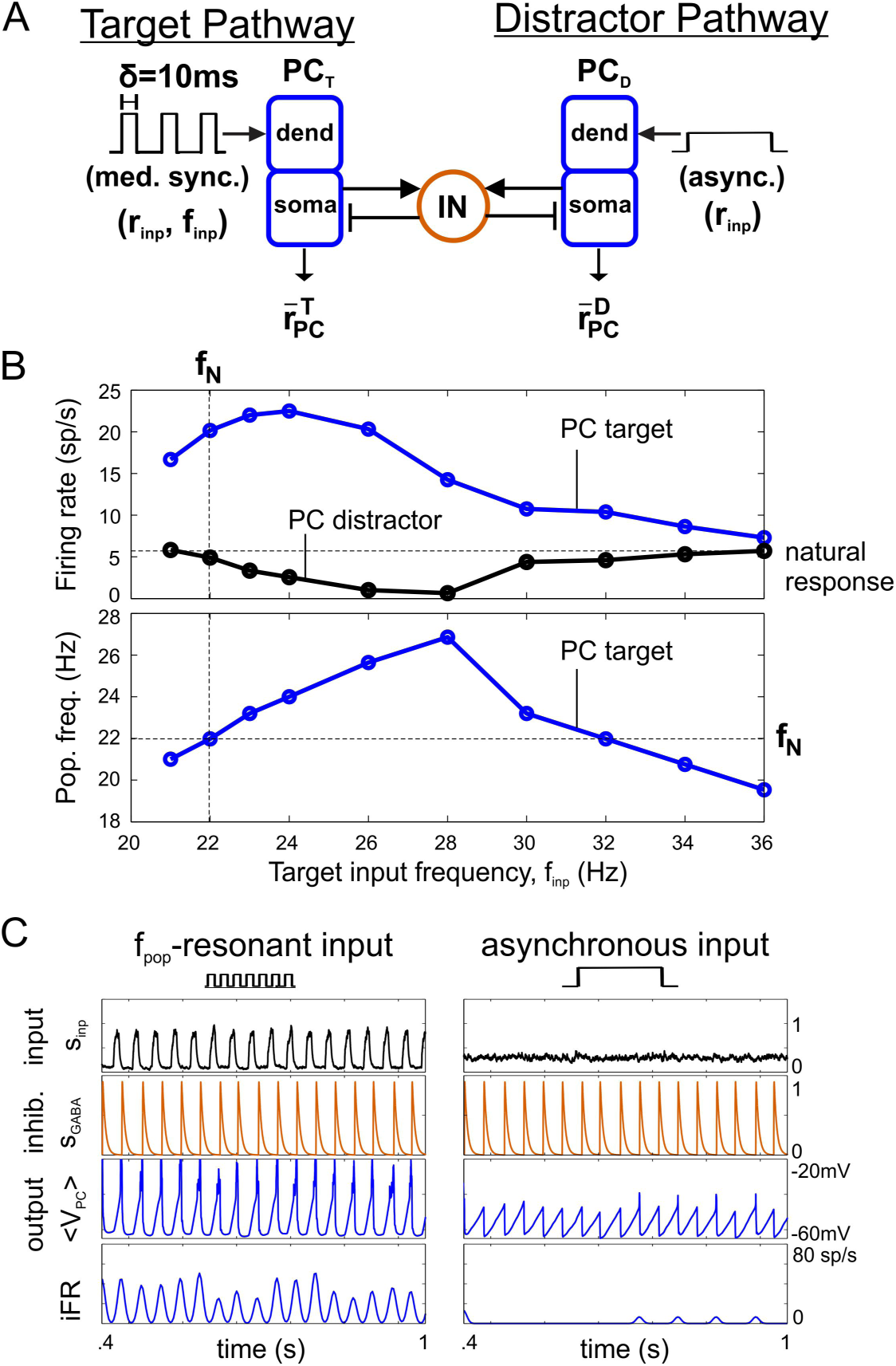
Frequency-dependent suppression of asynchronous activity. (**A**) Diagram showing a target PC population, PC_T_, driven by medium-synchrony oscillatory input in competition with an asynchronously-driven distractor PC population, PC_D_. (**B**) Dependence of distractor suppression on target input frequency. (i) Time-averaged firing rates (FRs) of PC_T_ (blue) and PC_D_ populations. As expected, PC_T_ FR peaks at the *r̄_PC_*-resonant frequency. PC_D_ responds at the FR expected given asynchronous input (horizontal black line, labeled “natural response”) when target input frequency is below the natural frequency (vertical black line) or far above 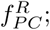 it is suppressed at intermediate frequencies. (ii) PC_T_ population (output) frequency versus the input frequency to PC_T_. As expected, PC_T_ *f_pop_* peaks at the *f*_pop_-resonant frequency. Importantly, whenever *f_pop_* exceeds the natural frequency (horizontal black line), PC_D_ FR is suppressed; maximal suppression of PC_D_ occurs when PC_T_ *f_pop_* is maximal and not when PC_T_ FR peaks in (i). (**C**) Example simulation with continuous suppression of the distractor pathway by a target pathway driven with a *f_pop_*-resonant input. On every cycle, the more rapidly oscillating target population engages the INs before the distractor reaches threshold.

Without competition, both PC populations would output periodic pulse packets of excitatory spikes, the target at the input frequency if 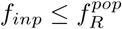 and the distractor at the natural frequency *f_N_*. The PC population frequency determines how frequently a PC population engages the IN population. When multiple outputs compete through shared INs, the output population with the highest frequency oscillation most frequently drives IN cells, tends to phase lock with them, and suppresses spiking in output populations oscillating more slowly. Any time the target population oscillates with a frequency faster than the natural frequency (i.e., *f_pop_* > *f_N_*), spiking in the distractor population is suppressed (Fig 3B). Importantly, peak suppression of the distractor population occurs when the target population frequency is maximal and not when the target PC activity is strongest. This implies that the optimal driving frequency to suppress responses to asynchronous distractors is the *f_pop_*-resonant frequency (Fig 3B-C). Such a rhythmically-driven output oscillating faster than the natural oscillation will always drive INs to continuously suppress responses to asynchronous distractors as long as the faster oscillation in the target remains. Internally-generated, nested oscillations with frequencies greater than *f_N_* would also suppress the response to asynchronous drive but only while present on the depolarizing phase of the slower driving oscillation. This distractor suppression occurs because the target population recruits interneuron-mediated lateral inhibition on every cycle of its oscillatory input with a period shorter than that required for the distractor population to reach threshold (Fig 3C, see membrane potential plots). Even if the lower-frequency distractor would otherwise have a higher firing rate than the target, its spike output is never fully realized when it receives another pulse of strong inhibition before reaching threshold. For this reason, the outcome of the competition is determined by the frequency of the population oscillation and not its amplitude. Dynamically, the suppression arises within a cycle as the target begins to oscillate more quickly than the distractor (S5 Fig). In contrast, there is no suppression of either pathway when the distractor input is strong enough so that the natural frequency that it induces equals the population frequency of the target (S6 Fig), despite the distractor PCs having lower firing rate, or when both PC populations receive asynchronous input (see T1 in S5 Fig).

Furthermore, the extent to which maximum 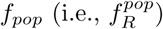 exceeds *f_N_* determines the range of input frequencies that can activate targets that suppress competing responses to asynchronous distractors (Fig 3B). Since 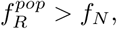 there is always an input frequency that can suppress competing distractors. In this scenario, the expected firing rate difference does not determine who is suppressive as long as firing rates are sufficient for a PC population pulse to activate the inhibitory INs. This result provides further justification for considering *f_pop_* as an output measure (in addition to *r̄_PC_*) because that frequency can determine the outcome of competition. This example represents a novel, functionally-relevant reason for examining output *f_pop_* and demonstrates the importance of the separation between the natural *f_N_* and resonant frequencies 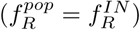 in PC/IN networks with strong feedback inhibition.

### Inputs tune the PC/IN network

For the remainder of the work presented here we will examine how the response properties of the PC/IN network (with one output PC population) depend on flexible parameters of the input and the slower effects of neuromodulation. It will be shown that the response properties of the PC/IN oscillator are not purely intrinsic and can be adaptively shaped by extrinsic influences. This represents a powerful means by which task-related modulations can influence cortical processing.

#### Input synchrony increases the separation between natural and peak frequencies

More synchronous input rhythms (i.e., smaller *δ_inp_*) delivering more coincident spikes (i.e., larger instantaneous drives) to each PC cell (Fig 4A) drove larger fractions of the target PC population to spike on each cycle before feedback inhibition was recruited to silence it. Consequently, greater synchrony enhanced output firing rates *r̄_PC_* for all input frequencies and the strength of resonant response (Fig 4Bi) without affecting the resonant input frequency 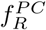 maximizing PC firing rates (Fig 4Bi, C). In contrast, 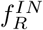 increased with input synchrony because *r̄_PC_* remained sufficiently large to engage interneurons for greater *f_inp_*; and, since 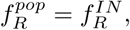 peak population frequency also increased (Fig 4C); this increase in separation between natural and peak frequencies with synchrony implies that a wider range of input frequencies can be exclusively selected (i.e., suppress responses to asynchronous activity) when they are more synchronous. In summary, output networks are able to achieve faster network oscillations, produce greater projection neuron output, and recruit more local inhibition when target signal inputs are more synchronous.

**Figure 4.**
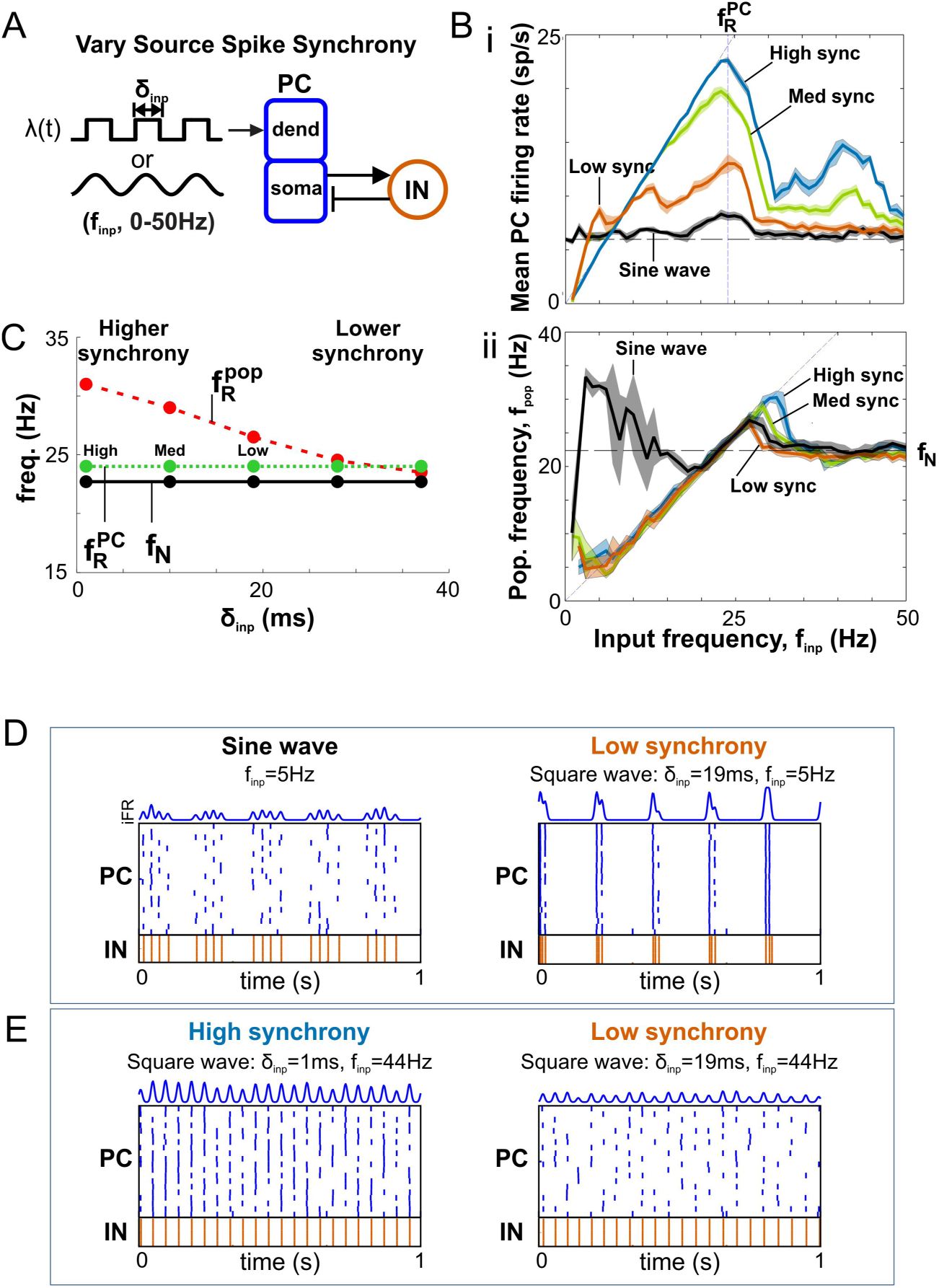
Dependence of response profiles on input synchrony. (**A**) Diagram of PFC network receiving variable-synchrony square wave or sinusoidal inputs. (**Bi**) Firing rate profile for PC populations given oscillatory inputs with different degrees of synchrony. (**Bii**) Population frequency profile for inputs with different degrees of synchrony. Horizontal dashed line marks the natural frequencies for each degree of synchrony. (**C**) The effect of input synchrony on resonant frequencies. Maximum population frequency (at the IN firing rate res. freq.) increases with input synchrony. (**D**) Spike rasters and PC iFR responses showing the nesting of natural oscillations generated by a local network on the depolarizing phase of a lower-frequency external driving oscillation with sine wave (left) or square wave (right) rate-modulation. (**E**) Spike rasters and PC iFR responses showing that weaker firing rate resonance at the first harmonic (i.e., smaller bump at *f_inp_* = 44Hz in Bi, blue curve) occurs for high synchrony (left) but not low synchrony (right) oscillatory inputs.

#### Input synchrony-dependent responses below the natural frequency and above the peak frequency

We discovered a number of noteworthy behaviors of the PC/IN network that depend on input synchrony at driving frequencies above and below the natural and resonant frequencies. When inputs have low synchrony, they can deliver a suprathreshold input to PCs that lasts longer than the duration of feedback inhibition, resulting in PC oscillations nested within each cycle of the input (Fig 4D). This represents a mechanism for generating nested oscillations through an interaction between a slow external driving rhythm (with low spike synchrony) and an internally generated, inhibition-based natural rhythm. These nested oscillations can produce second population frequencies that have more power than the input frequency (see the bump for low frequency sine wave inputs in Fig 4C and corresponding power spectra in S7 Fig) and mean rates that exceed the input frequency when PCs spike more than once per input cycle (see bumps at low frequencies in Fig 4B).

In contrast, PCs spike at most once per cycle when inputs are highly synchronous. Additionally, it is known that there is greater postsynaptic EPSP summation of more synchronous spikes. For highly synchronous inputs, this causes all PCs to spike on every cycle when there are enough input spikes driving them. Given square-wave inputs with a fixed number of total spikes (built-in to the study to achieve equal-strength rhythmic and asynchronous inputs), the number of spikes delivered per cycle decreases as frequency increases. This decrease in pulse strength with increasing frequency restricts the range of input frequencies that engage all PCs on every cycle. The dependencies of input pulse strength on synchrony and frequency cause the mean output rate to increase with input frequencies well below *f_N_* to an extent that scales with input synchrony (Fig 4B, compare low and high synchrony). Finally, well above the natural frequency, the *r̄_PC_* profile exhibited smaller peaks at harmonics of the resonant frequency in response to highly synchronous inputs (Fig 4E).

#### Stronger inputs increase natural and resonant frequencies

Stronger inputs (i.e., higher time-averaged rate *r_inp_*) (Fig 5A), delivering larger mean drives to each PC cell, increased the mean output firing rate (Fig 5Bi), natural and peak population frequencies (Fig 5Bii), and firing rate resonant frequencies (Fig 5C). The dependence of *f_N_* on *r_inp_* implies the natural response is a variable-frequency network oscillation controlled by the strength of input (see Discussion for functional implications). 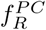 equaled *f_N_* for weak inputs and increasingly exceeded it for inputs with increasing strength; in contrast, 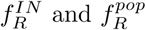 exceeded *f_N_* for all input strengths that were strong enough to produce a natural oscillation (Fig 5C). Finally, 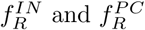 converged when the input was too weak to produce a natural oscillation and the network entered a band-pass regime (S1 Fig). The fact that the peak frequency always exceeds the natural frequency at drives where a natural oscillation is present implies that there is always an input frequency that enables suppression of responses to asynchronous activity.

**Figure 5.**
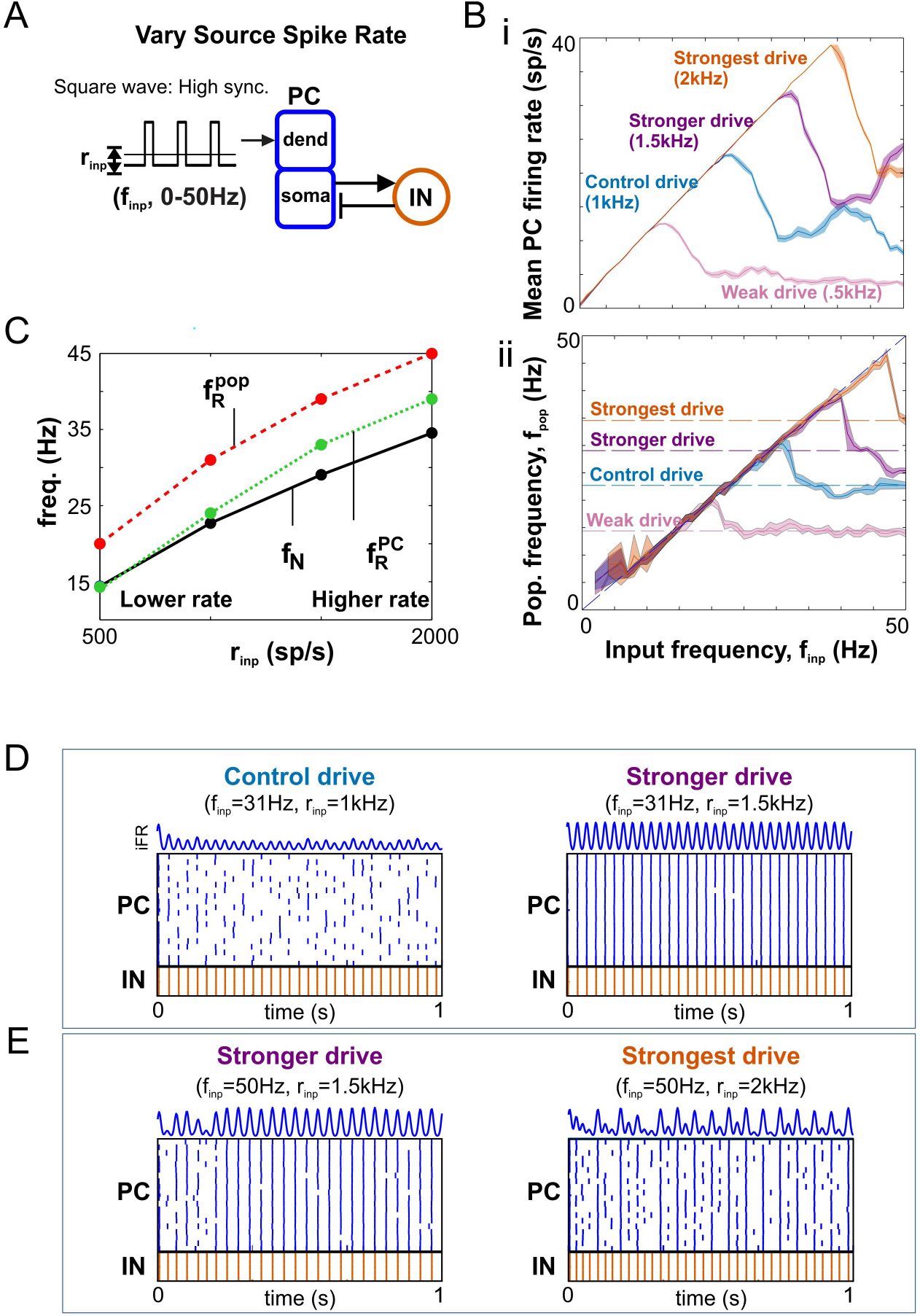
Dependence of response profiles on input strength. (**A**) Diagram of PFC network receiving variable-strength high-synchrony square wave input. (**Bi**) Firing rate profile for PC populations given oscillatory inputs with different strengths. (**Bii**) Population frequency profile for inputs with different strengths. Horizontal dashed lines mark the natural frequencies for each drive strength. (**C**) The effect of input strength on natural and resonant frequencies. (**D**) Spike rasters and PC iFR responses showing the typical case of stronger input driving more output: (left) weaker input, less output, (right) stronger input, more output. (**E**) Spike rasters and PC iFR responses showing special case of resonance at first harmonic enabling a weaker input to drive more output: (left) weaker input, more output, (right) stronger input, less output.

#### Neuromodulation of the PC/IN network

We have shown in previous sections that the control PC/IN network based on rat medial prefrontal cortex exhibits beta-range natural (Fig 1) and resonant (Fig 2) frequencies. Next, we simulated knockout experiments to explore how modulating the conductance of non-spiking currents would affect the network response (Fig 6). Removing hyperpolarizing currents (*I_Ks_*, *I_KCa_*) increased the *r̄_PC_*-resonant frequency, while removing depolarizing currents (*I_NaP_*) decreased the resonant frequency or (*I_Ca_*) silenced PCs altogether (Fig 6C). The weak effect of removing the hyperpolarizing currents could be amplified by increasing their conductance. As long as PCs remained in a spiking regime, removing modulatory currents did not qualitatively alter the response profile in most cases. The one exception was that removing *I_KCa_* resulted in a flatter profile near the peak; then across realizations, this caused resonant peaks to occur at neighboring input frequencies and produced a non-zero standard deviation on the 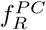 bar plot. A parsimonious explanation of these effects is that the shift in resonant frequency resulted from the shift in excitability caused by the non-spiking currents. Consistent with this hypothesis, similar shifts were achieved with the addition of tonic inputs that similarly shift baseline excitability (Fig 6D). Thus, neuromodulation of non-spiking currents and baseline excitability can tune the response profiles of PC/IN networks.

**Figure 6.**
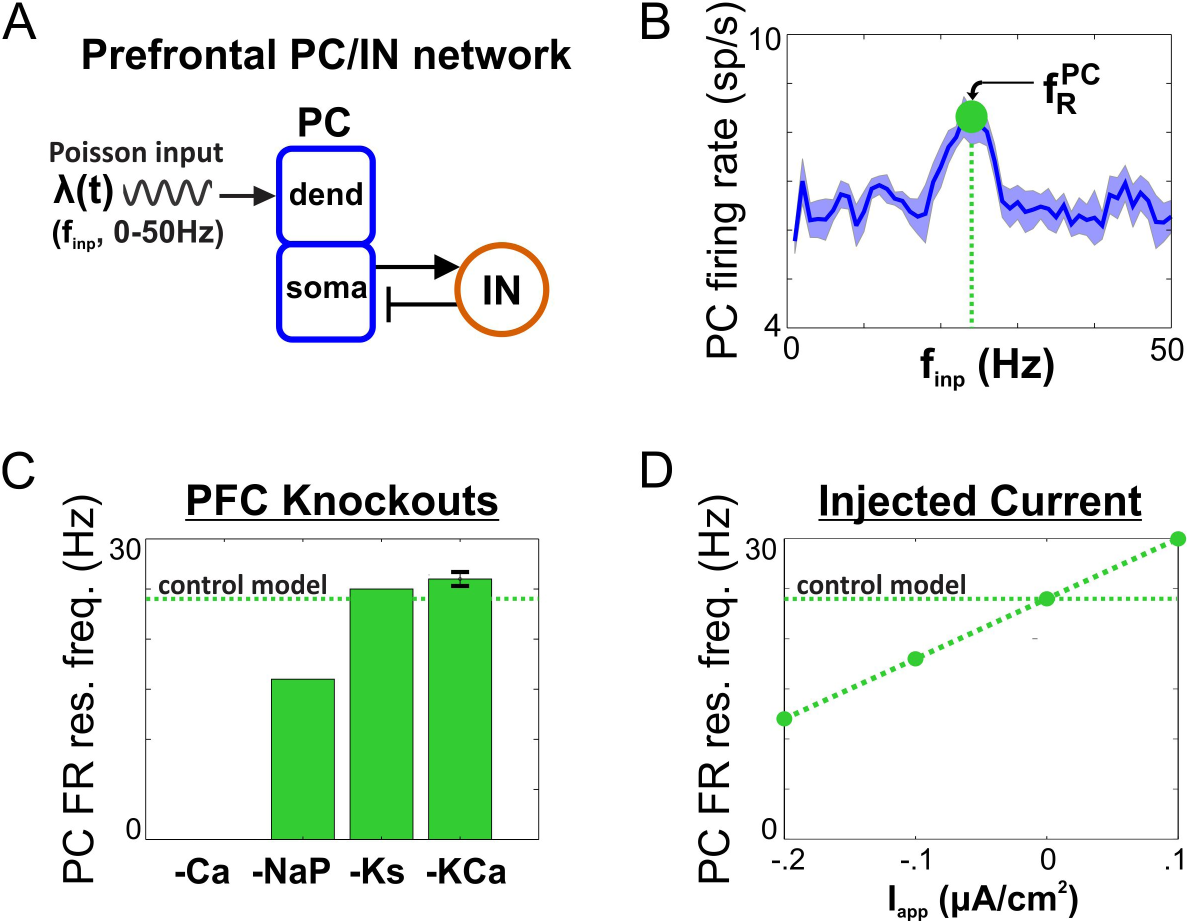
Neuromodulation of firing rate resonance in PC/IN network. (**A**) Diagram showing an external sinusoidal Poisson input to the dendrites of 20 two-compartment principal cells (PCs) receiving feedback inhibition from a population of 5 fast spiking interneurons (INs). PC and IN models include conductances found in prefrontal neurons (see Fig 7A for details). (**B**) Input frequency-dependent firing rate profile showing resonance at a beta2 frequency. (**C**) The effect of knocking out individual ion currents on the resonant input frequency maximizing firing rate outputs. Removing hyperpolarizing currents (-Ks, -KCa) increased the resonant frequency, while removing depolarizing currents (-NaP) decreased the resonant frequency or (-Ca) silenced the cell altogether (see Fig 7A for ion channel key). Error bars indicate mean ± standard deviation across 10 realizations; only -KCa had a non-zero standard deviation (i.e., values that differed across realizations). (**D**) The effect of hyperpolarizing and depolarizing injected currents, I_app_, on the resonant frequency mirrored the effect of knockouts on excitability.

**Figure 7.**
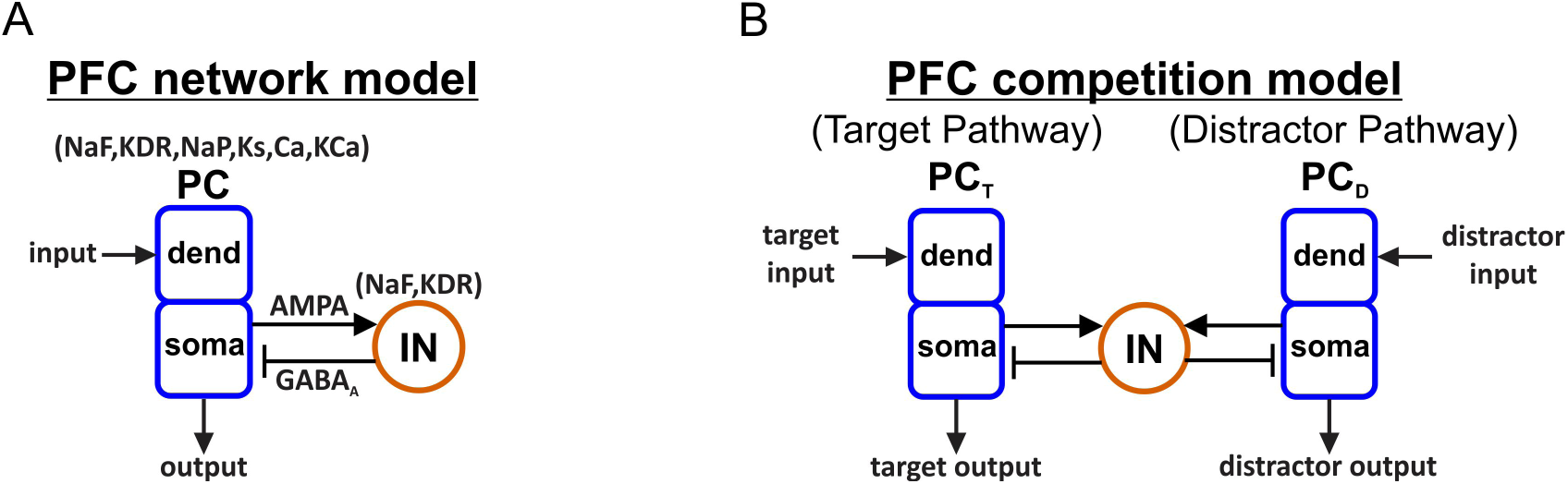
Architecture of output networks. (**A**) Diagram showing feedforward excitation from external independent Poisson spike trains to the dendrites of 20 two-compartment (soma, dend) principal cells (PCs) receiving feedback inhibition from a population of 5 fast spiking interneurons (INs). All PC and IN cells have biophysics based on rat prelimbic cortex (Ion channel key: NaF = fast sodium channel; KDR = fast delayed rectifier potassium channel; NaP = persistent sodium channel; Ks = slow (M-type) potassium channel; Ca = high-threshold calcium channel; KCa = calcium-dependent potassium channel). (**B**) Diagram showing a rhythmically-driven target population of PC cells (PC_T_) competing with an asynchronously-driven distractor population (PC_D_) through a shared population of inhibitory IN cells.

## Discussion

In this work, we characterized the prefrontal PC/IN network response to strong oscillatory inputs in terms of biologically-relevant input and output properties. The PC/IN network with strong feedback inhibition exhibited resonance in the spiking of PC and IN populations as well as the output population frequency of the network. We have shown that a separation of preferred frequency for output spiking (the frequency maximizing PC activity) and the maximal frequency that can be relayed by the network is enabled by the combination of (1) strong excitatory input that generates a response to all input frequencies, (2) strong feedback inhibition: the ability of fast spiking INs to synchronously silence the PC population, and (3) noisy spiking in a population of PCs for which an active subset are able to activate INs. The peak output frequency of the inhibition-based network oscillator was determined by the input frequency maximizing local inhibition from the INs and always exceeded the natural frequency induced by equal-strength asynchronous activity. This population frequency resonance was shown to determine the optimal driving frequency for suppressing responses to asynchronous input. Finally, we showed that the resonant properties of PC/IN networks can be flexibly tuned by task-modulated signal properties (synchrony and strength) to dynamically shape ongoing neural processing.

### Functional implications

#### Boosting: *Amplifying signals with preferred frequencies*

Firing rate resonance (i.e., *r̄_PC_*-resonance) in neuronal networks can be used to amplify population signals embedded in a resonant oscillation. Such amplification has been shown to promote the propagation of signals across weakly connected brain areas [19] and to support the transmission of time-varying, rate-coded signals when signal fluctuations are slow relative to the resonant frequency [10]. The smaller *r̄_PC_*-resonances we observed at higher harmonics could enable synchronous signals carried at higher frequencies to benefit from the same effects.

High beta-frequency (20-35Hz) oscillations have been observed in prefrontal cortex (PFC) in numerous studies [4,14,20]. Here, we have shown that a PC/IN network constrained by prefrontal data exhibits *r̄_PC_*-resonance in the same range for a wide variety of inputs (i.e., oscillatory inputs with firing rates and synchrony levels spanning those observed experimentally in the same region). This beta resonance suggests that PFC networks are tuned for processing signals embedded in beta rhythms. Furthermore, we have shown that the natural response of the prefrontal network driven by asynchronous spiking is to generate beta-frequency oscillations. This could explain why beta rhythms are frequently associated with top-down cognitive control of attention [21,22] and decision making [23,24].

Transcranial stimulation is often used to enhance neural oscillations [25,26]. [27] showed that transcranial alternating current stimulation (tACS) with sawtooth waves is more effective at enhancing alpha-frequency oscillations than tACS with sinusoidal waves; whether tACS predominantly excites interneurons or principal cells depends on the intensity of stimulation [28]. Our work suggests that, given an excitatory intensity, tACS stimulation with square waves (i.e., periodic pulses) could be even more effective at enhancing neural oscillations. Furthermore, the relationship between natural and resonant frequencies suggests an experimental protocol for maximally activating a region using a fixed excitatory intensity: first, apply a continuous pulse of direct current stimulation (tDCS) while recording EEG to identify the natural frequency of a target region; then, use equal-intensity tACS at the same or slightly higher frequency. This approach would enable maximal activation of a region near its preferred frequency following a single direct current stimulation. It also provides more specific activation of target regions with corresponding resonant properties. The protocol could be validated experimentally by comparing the tDCS response to a set of tACS responses with different stimulation frequencies. The natural frequency could be computed as the beta/gamma frequency (i.e., potential frequencies for inhibition-paced network oscillators) with peak EEG power following tDCS, while the resonant frequency is the tACS stimulation frequency maximizing EEG power around the stimulation frequency. The same protocol could be performed using transcranial magnetic stimulation (TMS) and rhythmic TMS.

#### Gating: *Selecting outputs based on preferred frequencies*

We have also shown that, while peak PC firing determines maximum spike output in a network with one PC population, it is population frequency that determines whether responses to asynchronous activity will be suppressed in PC populations competing through IN-mediated lateral inhibition. This enables exclusive response to oscillatory inputs, demonstrated in this work with one PC population driven by an *f_pop_*-maximizing oscillatory input and the other by an asynchronous input (Fig 3). It is the output population oscillating faster in response to external oscillations that dominates control of the interneuron population from cycle to cycle and which effectively suppresses the opposing response to asynchronous drive.

Asynchronous activity does not necessarily reflect background noise [29]; whether it is signal or noise, our work suggests resonant oscillatory activity may be given priority over it. PFC models of working memory (WM) often include item representations maintained in asynchronous, persistent activity [30, 31], and a similar encoding state has been implemented in neuromorphic hardware [32]. However, working memory items have also been found to be phase-locked to high-frequency beta rhythms [14], and spiking models of WM items in oscillatory states have been developed [31,33,34]. We hypothesize that WM representations, maintained in superficial layers of PFC [35], can encode items in both asynchronous and oscillatory states, and that the latter are given priority due to the resonant properties of the deep output layer that we investigated in this work. As we have shown, a deep layer PC population driven by a *f_pop_*-resonant oscillatory item in a superficial WM buffer would suppress the response in PC populations driven by items in an asynchronous state.

In the PC/IN network, increasing feedforward inhibition decreases the PC response to asynchronous and oscillatory inputs. For sufficiently strong feedforward inhibition, the PC/IN network is transformed into a bandpass filter that responds exclusively to sinusoidal inputs with a frequency near the *r̄_PC_*-resonant frequency of the network (see Methods for a comparison of filter properties given sine vs. square wave inputs); in this case, asynchronous inputs have no effect on the PC population. [10] used an IN network with firing rate resonance and sinusoidal input to deliver bandpass-creating feedforward inhibition to a PC population; in their case, the bandpass PC response depended further on a phase lag between the excitatory input and feedforward inhibition at the *r̄_IN_*-resonant frequency. With this setup, they show how bandpass filters can be used to de-multiplex target signals from a mixture of converging inputs. The prefrontal PC/IN network could similarly de-multiplex signals given bandpass-creating feedforward inhibition and sinusoidal input.

#### Modulation: *Tuning the output and preferred input frequencies*

[36] showed that the resonant frequency can be tuned by changing the connection weights among excitatory and inhibitory populations (i.e., “rewiring the network topology”). Here, we show that the resonant and natural frequencies of similar networks can be tuned dynamically by changing the strength of an oscillatory input or the baseline excitation in the output PC population. The latter can be tuned through neuromodulation (Fig 6C) (e.g., modulation of potassium currents in PFC by dopamine [37] and acetylcholine [38]) or external applied currents (Fig 6D) representing modulatory signals with asynchronous spiking.

One consequence of the dependence of the natural output frequency on input firing rate (Fig 5C) is that PC/IN networks with strong feedback inhibition can operate as variable-frequency oscillators. If a PC/IN network outputs to a bank of band-pass filter networks with different center frequences, this could enable input rate-based control of which filter network is activated. In this scenario, a PC/IN network driven by asynchronous spiking would effectively perform a firing rate-to-oscillation frequency conversion that could be used to route signals to select elements of a downstream filter bank. The fact that the output frequency depends on the time constant of feedback inhibition means that correspondence between output and center frequencies could be facilitated by matching interneuron types in the converter and filter networks. Furthermore, the variable-frequency response could potentially serve encoding of slowly-varying asynchronous signals using pulse-frequency modulation [39] or participation in a phase-locked loop. Such systems would need to account for, or be invariant to, the concurrent amplitude modulation of the PC/IN network. The fact that the output rhythms are sparse (i.e., only a fraction of PCs spike on every cycle) makes the signal energy efficient and potentially suitable for use in neuromorphic engineering [40,41].

Together, these results demonstrate flexibility of neural processing provided by extrinsic tuning of PC/IN oscillator properties.

### Relation to other work

Resonance phenomena have been studied in neural systems at multiple scales. Peaks in the single neuron membrane potential response to subthreshold oscillatory inputs have been studied in terms of the interplay between intrinsic ion currents [9]; their ability to influence spiking has been demonstrated in single neurons [42]; and relationships between subthreshold resonance and the natural network frequency of electrically coupled excitatory cells have been shown [43]. Our PC model, in isolation, exhibits subthreshold resonance at delta frequencies ( 2Hz) (S1 Fig A) that translates into an input strength-independent spiking resonance at the same frequency for suprathreshold inputs (S1 Fig B-C). The addition of strong feedback inhibition suppresses the spiking response to delta-frequency inputs while a higher-frequency, input strength-dependent spiking resonance emerges in response to strong inputs (S1 Fig D). We have explored this higher-frequency resonance in this work and shown how it depends on the strength of input (S1 Fig Ei-ii) and the time constant of feedback inhibition (S1 Fig Eiii).

The mechanism that determines the precise value of 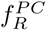 in the inhibition-based PC/IN oscillator is not fully understood. Insight into the locking of an inhibition-based IN network oscillator to periodic drive with heterogeneous phases has been obtained using a timing map [18]; a similar analysis might provide insight in the present case of a PC/IN network locking preferentially to a particular periodic drive with heterogeneous spike times but is beyond the scope of this work. The work in [18] and our results suggest the value of 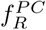 is related to the number of PC spikes that are necessary to engage INs on a given cycle. Changing the network size while keeping the synaptic strengths constant did not affect 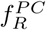 (compare 2Bi to S4 Fig). This is in contrast to work showing that resonant frequency in a network model of Wilson-Cowan oscillators depended strongly on network size [44]. Further work is needed to understand the differences between resonances in spiking versus activity-based networks.

[11] showed that strong feedback inhibition is required for firing rate resonance in integrate-and-fire networks of excitatory and inhibitory cells driven by sinusoidal inputs; they also showed that stronger inputs increase resonant frequencies. We have reproduced these qualitative findings in a more detailed network model and extended the results by showing how firing rate resonance relates to the natural and peak oscillation frequencies in the strong-input regime, and how they all depend on other properties of the oscillatory input (i.e., waveform and spike synchrony). Compared to the integrate-and-fire networks (S3 Fig), the prefrontal network exhibited lower natural and resonant frequencies. [45] showed that heterogeneity of PC intrinsic properties in a PC/IN network produces a range of resonant frequencies that supports combining, instead of selecting, inputs. In contrast, our PC population is homogeneous, and the network produces more selective responses favoring outputs with *f_pop_*-resonant inputs. See Methods for a comparison of output measures used in this and other studies of spiking resonance. [46] investigated the response of an inhibition-paced IN population to physiologically-relevant periodic pulse inputs, but their work did not include PC cells or examine resonance.

### Limitations and future directions

The model investigated in this work made the following simplifications: no NMDA synapses, usage of all-to-all connectivity between PCs and INs, no PC-to-PC or IN-to-IN connectivity, and the lack of noise driving INs. Preliminary simulations demonstrated that probabilistic connectivity, weak noise, and weak NMDA synapses did not disrupt the results. For strong noise to INs, additional IN-to-IN feedback inhibition is necessary to synchronize the IN population. Furthermore, IN-to-IN and PC-to-PC connectivity have been shown to modulate resonant frequencies [36]. We suspect the main requirement for our results to hold is that the network is in a regime that produces a natural oscillation in response to an asynchronous input to the PCs (i.e., external noise to INs must be weak enough, feedback inhibition strong enough, and PC-to-IN drives strong and fast enough for a fraction of spiking PC cells to control IN activity from cycle to cycle).

Another important limitation of the present work pertains to the role of modulatory intrinsic currents. We have focused on regimes where PCs are roughly regular spiking and INs are fast spiking. More work is needed to understand how the dynamics reported here would be affected by PCs that are intrinsically bursting (as observed in deep layers of cortex and thalamus) and INs exhibiting low-threshold spiking. Furthermore, our account of the effects of knocking out modulatory currents is limited to effects on overall activity levels across the PC population. In contrast, work by [47] shows that modulatory currents can impact the cycle-to-cycle probability of individual cells participating in the population rhythm.

### Conclusions

The work reported here has introduced a distinction between time-averaged firing rate resonance in excitatory PCs and inhibitory INs that arises when inputs are strong. We have also introduced a new form of network resonance observed in strongly-driven inhibition-based PC/IN oscillators, called population frequency resonance that depends on firing rate resonance in INs; and demonstrated its importance for suppressing responses to asynchronous activity. These results have also made a significant contribution to understanding how PC/IN networks with strong feedback inhibition are affected by task-modulated changes in oscillatory inputs, in general, and why beta rhythms are so frequently associated with prefrontal activity.

## Materials and methods

### Network models

The network model represents a cortical output layer with 20 excitatory principal cells (PCs) connected reciprocally to 5 inhibitory interneurons (INs). Hodgkin-Huxley (HH) type PC and IN models were taken from a computational representation of a deep layer PFC network consisting of two-compartment PCs (soma and dendrite) with ion channels producing *I_NaF_*, *I_KDR_*, *I_NaP_*, *I_Ks_*, *I_Ca_*, and *I_KCa_* currents (μA/cm^2^) and fast spiking INs with channels producing *I_NaF_* and *I_KDR_* currents [48] (Fig 7A; see figure caption for channel definitions). IN cells had spike-generating *I_NaF_* and *I_KDR_* currents with more hyperpolarized kinetics and faster sodium inactivation than PC cells, resulting in a more excitable interneuron with fast spiking behavior [48]. In the control case, PC and IN cell models were identical to those in the original published work [48] while network connectivity was adjusted to produce natural oscillations (not in [48]), as described below, and the number of cells in the network was decreased to enable exploration of larger regions of parameter space while remaining large enough to capture the dynamics of interest for this study; however, the same resonant frequencies were obtained in simulations using the original network size (S4 Fig). Knockout experiments were simulated by removing intrinsic currents one at a time from the PC cell model. All cells were modeled using a conductance-based framework with passive and active electrical properties of the soma and dendrite constrained by experimental considerations [49]. Membrane potential *V* (mV) was governed by:

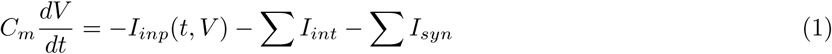

where *t* is time (ms), *C_m_* = 1 μF/cm^2^ is the membrane capacitance, *I_int_* denotes the intrinsic membrane currents (μA/cm^2^) listed above, *I_inp_*(*t*, *V*) is an excitatory current (μA/cm^2^) reflecting inputs from external sources described below, and *I_syn_* denotes synaptic currents (μA/cm^2^) driven by PC and IN cells in the network. We chose to explore the prefrontal model as part of a larger project on prefrontal oscillations. We confirmed the generality of our qualitative results using a leaky integrate-and-fire (LIF) model; see the caption of S3 Fig for details on the LIF model. We explored single cell and minimal network versions of our HH type model to investigate potential relationships between single cell and network resonances; details on these simulations can be found in the caption of S1 Fig.

The output layer had either one or two populations of PC cells with each output population receiving either one or two input signals. Input frequency-dependent response profiles were characterized using a network with one input and one output (Fig 7A). Competition between the outputs of parallel pathways was investigated using a network with two homogeneous output populations receiving one input each while interacting through a shared population of inhibitory cells (Fig 7B).

### Network connectivity

PC cells provided excitation to all IN cells, mediated by *α*-amino-3-hydroxy-5-methyl-4-isoxazolepropionic acid (AMPA) currents. IN cells in turn provided strong feedback inhibition mediated by *γ*-aminobutyric acid (GABA_A_) currents to all PC cells. This combination of fast excitation and strong feedback inhibition is known to generate robust network oscillations in response to tonic drive [12,13]. AMPA currents were modelled by:

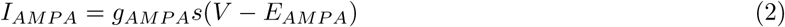

where *V* is the postsynaptic membrane voltage, *g_AMPA_* is the maximal synaptic conductance, *s* is a synaptic gating variable, and *E_AMPA_* = 0 mV is the synaptic reversal potential. Synaptic gating was modeled using a first-order kinetics scheme:

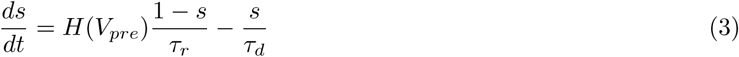

where *V_pre_* is the presynaptic membrane voltage, *τ_r_* = 0.4 ms and *τ_d_* = 2 ms are time constants for neurotransmitter release and decay, respectively, and *H*(*V*) = 1 + *tanh*(*V*/4) is a sigmoidal approximation to the Heaviside step function. GABA_A_ currents are modeled in the same way with *E_QABA_* = –75 mV and *τ_d_* = 5 ms. Maximum synaptic conductances for PC cells were (in mS/cm^2^): GABA_A_ (.1); for IN cells: AMPA (1).

### External inputs

Each PC cell received independent Poisson spike trains (Fig 8) with time-varying instantaneous rate λ(*t*) (sp/s) and time-averaged rate *r_inp_* = 〈λ〉; spikes were integrated in a synaptic gate *s_inp_* with exponential AMPAergic decay contributing to an excitatory synaptic current *I_inp_* = *g_inp_s_inp_*(*V* – *E_AMPA_*) with maximal conductance *g_inp_* (mS/cm^2^). Input signals were modeled by collections of spike trains with the same instantaneous rate-modulation. A given input signal to a PC output population can be interpreted as conveying rate-coded information from a source population in a particular dynamical state.

**Figure 8.**
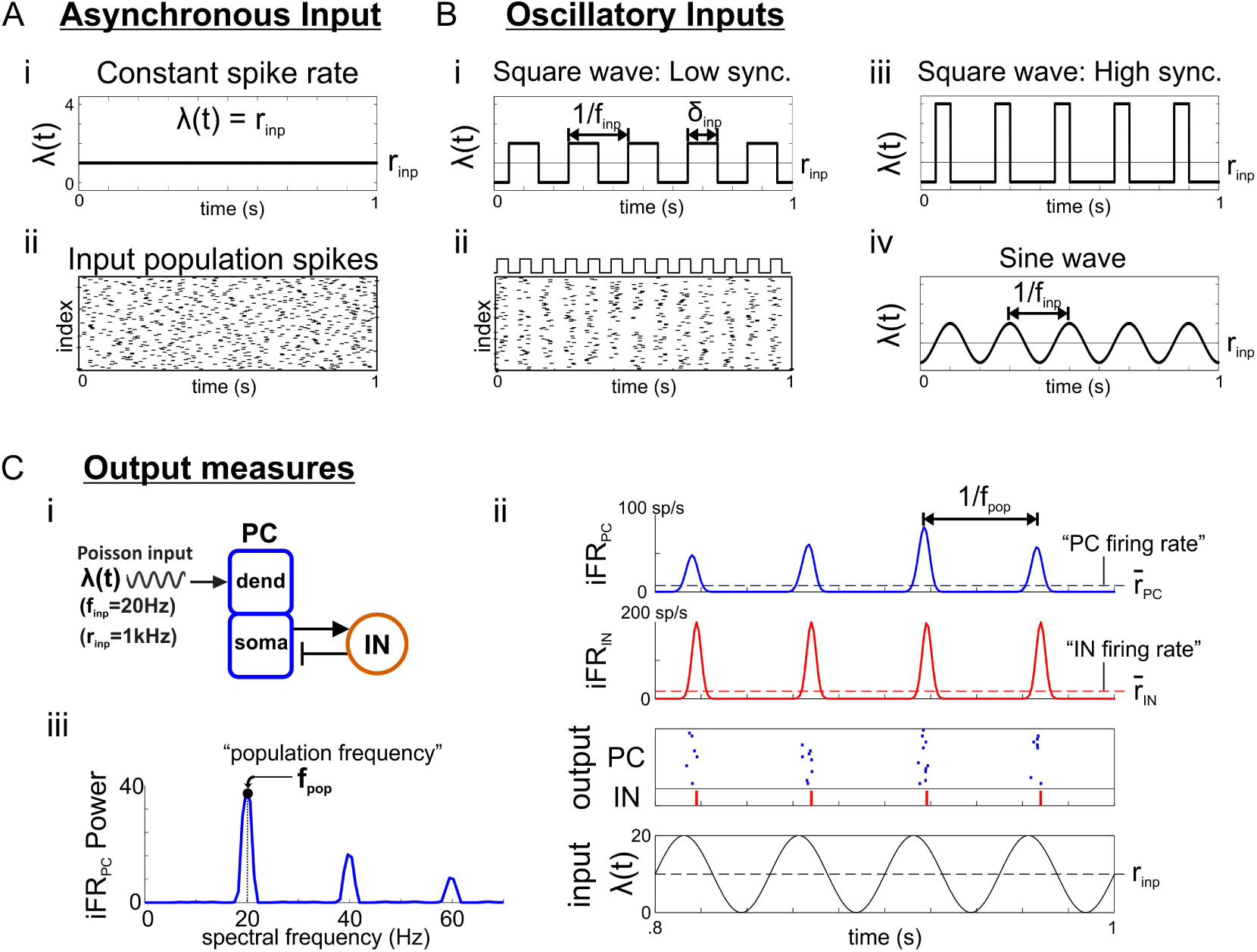
Input network activity. (**A**) Asynchronous Poisson input with (i) constant instantaneous rate *r_inp_* and(ii) raster for 100 input cells with *r_inp_* = 10 sp/s (equivalent to 1 input cell with *r_inp_* = 1000 sp/s). (**B**) Poisson inputs with oscillatory instantaneous rate-modulation. (i) Instantaneous rate modulated by low synchrony square wave, parameterized by pulse width *δ_inp_* and inter-pulse frequency *f_inp_*. (ii) raster plot produced by square wave input. (iii) High synchrony, square wave rate-modulation. (iv) sine wave modulation, parameterized by frequency *f_inp_*. (**C**) Output measures for the PC/IN network. (i) Diagram of a PC/IN network receiving an input from (B). (ii) Plots showing the instantaneous firing rate (iFR) computed for each population using Gaussian kernel regression on the spike raster. Time-averaged firing rates are defined by the mean iFR for each population. (iii) Power spectrum showing how population frequency is defined by the spectral frequency with peak power in the iFR.

Signals from sources in different dynamical states were generated by modulating instantaneous rates λ(*t*) according to the activity patterns exhibited by populations in those states. Signals from source populations in an asynchronous state were modeled by Poisson spike trains with constant rate λ(*t*) = *r_inp_* (Fig 8A) whereas signals from sources in an oscillatory state were modeled using periodically-modulated instantaneous rates (Fig 8B). Signals with sine wave modulation had λ(*t*) = *r_inp_* (1 + *sin*(2*π f_inp_t*)) /2 parameterized by *r_inp_* (sp/s) and rate modulation frequency *f_inp_* (Hz). Sinusoidal modulation causes spike synchrony (the interval over which spikes are spread within each cycle) to covary with frequency as the same number of spikes become spread over a larger period as frequency decreases. Thus, we also investigated oscillatory inputs with square wave modulation in order to differentiate the effects of synchrony and frequency while maintaining the ability to compare our results with other work. Square wave rate-modulation results in periodic trains of spikes with fixed synchrony (pulse packets) parameterized by *r_inp_* (sp/s), inter-pulse frequency *f_inp_* (Hz), and pulse width *δ_inp_* (ms). *δ_inp_* reflects the synchrony of spikes in the source population with smaller values implying greater synchrony; decreasing *δ_inp_* corresponds to decreasing the duty cycle of the square wave. For the square wave input, we chose to hold constant *r_inp_* so that across frequencies the only significant change is in the patterning of spikes and not the total number of spikes; this results in larger pulses being delivered to postsynaptic PCs at lower frequencies as would be expected if lower frequencies are produced by larger networks [50]. If the number of spikes per cycle was fixed, instead, as would be the case for a given input population with iFR fluctuating more rapidly and all cells spiking on every cycle, then the mean strength of the input would increase with frequency, and its effect on resonance would no longer be comparable to a sinusoidal input with increasing frequency. The consequence of holding the number of spikes per cycle fixed for a square wave input is discussed further below and related to the results for fixed-mean square waves in S8 Fig.

All principal cells in the output layer received additional asynchronous inputs representing uncorrelated background activity from 100 cells in other brain areas spiking at 1 sp/s. Notably feedforward inhibition was excluded from the present work so that asynchronous inputs were maximally effective at driving PC cells. The effects of adding feedforward inhibition and conditions under which each case holds are considered in the Discussion. Control values for input parameters were *r_inp_* = 1000 sp/s (corresponding to a source population with 1000 projection neurons spiking at 1 sp/s); *δ_inp_* = 1 ms (high synchrony), 10 ms (medium synchrony), or 19 ms (low synchrony), and *g_inp_* = .0015 mS/cm^2^. High synchrony inputs are similar to strong, periodic spikes while medium and low synchrony inputs distribute spikes uniformly over intervals comparable to sine waves at 100 Hz and 53 Hz, respectively.

In simulations probing resonant properties, the input modulation frequency *f_inp_* was varied from 1 Hz to 50 Hz (in 1 Hz steps) across simulations. In simulations exploring output gating among parallel pathways, input signals had the same mean strength (i.e., *r_inp_*); this ensures that any difference between the ability of inputs to drive their targets resulted from differences in the dynamical states of the source populations and not differences in their activity levels.

### Data analysis

For each simulation, instantaneous output firing rates, iFR, were computed with Gaussian kernel regression on population spike times using a kernel with 6 ms width for visualization and 2 ms for calculating the power spectrum. Mean population firing rates, *r_PC_* and *r_IN_*, were computed by averaging iFR over time for PC and IN populations, respectively; they index overall activity levels by the average firing rate of the average cell in the population (Fig 8Ci-ii). The frequency of an output population oscillation, *f_pop_*, is the dominant frequency of the iFR oscillation and was identified as the spectral frequency with peak power in Welch’s spectrum of the iFR (Fig 8Ciii, S7 Fig). As defined, *f_pop_* usually reflects the rhythmicity of internal spiking; however, when nested oscillations are present at low frequencies, *f_pop_* may reflect either internal or external rhythm frequencies (see Fig 4D for a raster plot and PC iFR, Fig 4B for a *f_pop_* response profile, and S7 Fig for an iFR power spectrum with nested oscillations). This ambiguity does not interfere with our study of resonance at higher frequencies where the signal has a single dominant frequency; however, a disambiguating measure of population frequency would be necessary to study regimes in which multiple frequencies are strongly present (e.g., strong, low-frequency, low-synchrony periodic inputs). The natural frequency *f_N_* of the output network was identified as the population frequency *f_pop_* produced in response to an asynchronous input.

Our measure of spiking activity in the strongly-driven network differs from measures used in work on resonance in weakly-driven networks [10,11,42]. In the weakly-driven (i.e., linear) regime, the iFR amplitude scales linearly with the input and can serve as a measure for detecting resonance. However, in the strongly-driven regime that we explore, iFR may scale nonlinearly with the input; in the case of high-synchrony inputs, iFR amplitude saturates below the resonant frequency (i.e., all cells spike once on every cycle), and it has a more complicated profile and scaling with input strength in other cases. [51] has explored spiking resonance in a strongly-driven single cell and defined a measure called spike frequency that is roughly equivalent to the time-averaged firing rate. We have chosen to use a similar measure, the time-averaged iFR, *r*_∗_, to capture overall increases or decreases in the amount of spiking produced in the strongly-driven network.

Qualitatively, *r_PC_* profiles differ for the PFC PC/IN network with strong feedback inhibition depending on the waveform of the periodic input (Fig 9). Weak sinusoidal inputs produce band-pass responses like those observed in [10,11] (Fig 9Ai; S1 Fig D-Ei, blue curve). Increasing the strength of those inputs produces an all-pass regime in which inputs at all frequencies elicit a response, although a resonant peak remains (Fig 9Aii; S1 Fig D-Ei, black curve). In contrast, a weak square wave with mean input held constant across frequencies produces a low-pass response due to the larger input pulses at low frequencies (Fig 9Bi). However, the curve still exhibits a peak that occurs at the same input frequency as for the sine wave given equal-strength input. Finally, a weak square wave with pulse amplitude held constant produces a high-pass response due to the increasing input strength that occurs with an increasing number of pulses (Fig 9Ci). Increasing the strength of square wave inputs also moves the network into an all-pass regime (Fig 9Bii, Cii), but only the fixed-mean square wave exhibits a resonant peak (Fig 9Bii). In this work, we focus on the sine and square wave cases where mean input strength is held fixed and resonance is well-defined in physiologically-relevant frequency ranges.

**Figure 9:**
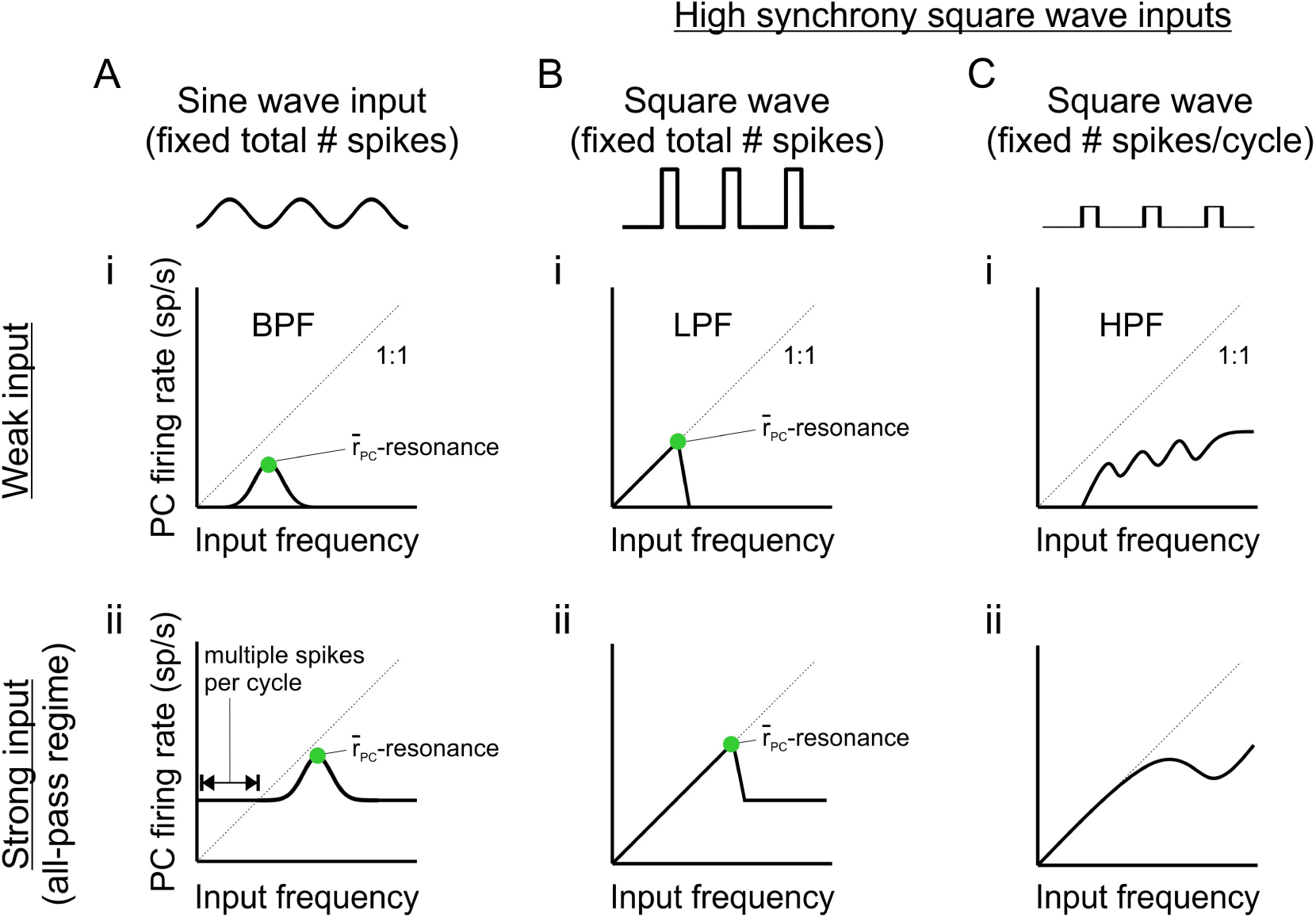
Cartoon profiles of time- and population-averaged PC firing rates in response to different types of oscillatory inputs. (**A**) Response to sinusoidal drive. (i) Weak input produces a band-pass filter (BPF) response with spikes driven by near-resonant frequencies and only a fraction of cells spiking on every cycle. (ii) Strong input produces an all-pass response with a resonant peak. **B**) Response to fixed-mean square wave drive. (i) Weak input produces a low-pass filter (LPF) response with spikes driven by all frequencies below a resonant peak and all cells spiking on every cycle. (**C**) Response to fixed-amplitude square wave drive. (i) Weak input produces a high-pass filter (HPF) response. (ii) Strong input produces an all-pass response without a well-defined resonant peak.

Across simulations varying input frequencies, statistics were plotted as the mean ± standard deviation calculated across 10 realizations. Input frequency-dependent plots of mean firing rates and population frequencies will be called response profiles. The time-averaged firing rate resonant frequencies,
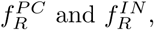
are the input frequencies at which global maxima occurred in the and *r_PC_* and *r_IN_* firing rate profiles, respectively. Similarly, the resonant input frequency,
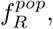
maximizing output oscillation frequency was found from peaks in *f_pop_* population frequency profiles, excluding the peaks that are due to nested oscillations in response to strong, low-frequency, low-synchrony periodic drives.

### Simulation tools

All models were implemented in Matlab using the DynaSim toolbox [52] (http://dynasimtoolbox.org) and are publicly available online at: http://github.com/jsherfey/PFC_models. Numerical integration was performed using a 4th-order Runge-Kutta method with a fixed time step of 0.01 ms. Simulations were run for 2500ms and repeated 10 times. The network was allowed to settle to steady-state before external signals were delivered at 400 ms. Plots of instantaneous responses begin at signal onset. The first 500 ms of response was excluded from analysis, although including the transient did not alter our results significantly.

## Supporting Information

**S1 Fig.**
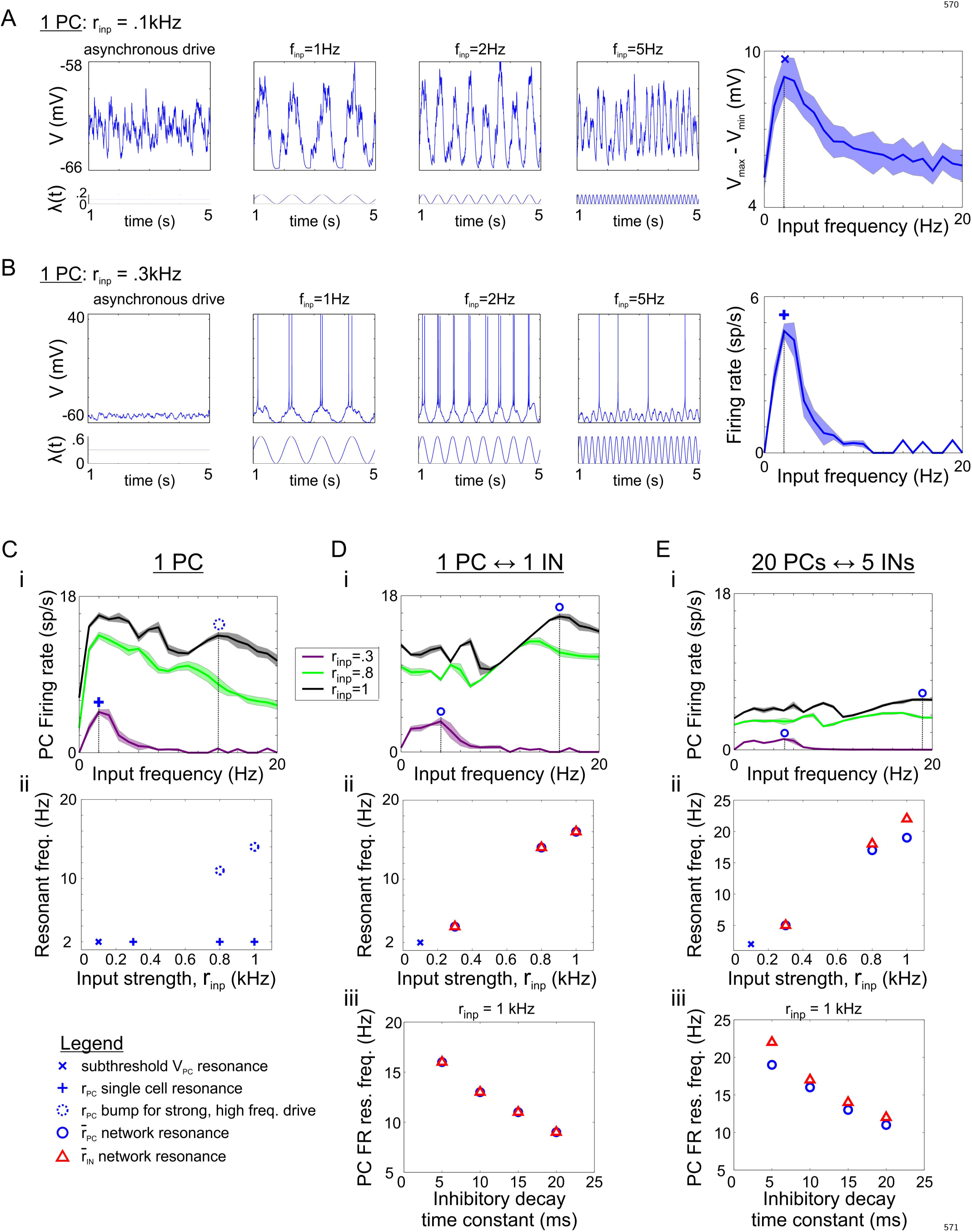
Comparison of types of resonance in the single PC and PC/IN networks. (**A**) After removing background noise to allow for subthreshold fluctuations, subthreshold resonance in the voltage fluctuation was observed at 2 Hz in a single principal cell (PC) driven by a weak sinusoidal drive (*r_inp_* = .1 kHz). Example voltage traces are shown in response to asynchronous input and sinusoidal inputs at f = 1 Hz, 2 Hz, and 5 Hz. The amplitude of voltage fluctuation, *V_max_* – *V_min_* is plotted versus input frequency, and the peak is marked with a × symbol. (**B**) After the input strength was increased to a slightly suprathreshold level (*r_inp_* = .1 kHz), suprathreshold spiking resonance was observed at the same 2 Hz frequency in a single PC. Example voltage traces are shown in response to asynchronous input and sinusoidal inputs at f = 1 Hz, 2 Hz, and 5 Hz. The time-averaged firing rate (FR) is plotted versus input frequency, and the peak is marked with a + symbol. This suggests that the subthreshold resonance translates to suprathreshold resonance in the linear regime. (**C**) (i) FR profile showing that as the strength of sinusoidal input is increased further to *r_inp_* = .8 kHz and 1 kHz, spiking resonance in the single PC remains at the same frequency and multiple bumps in time-averaged firing rate emerge at higher frequencies. The dotted circle marks the frequency at which an input strength-dependent bump in firing rate occurs (in the single PC) that is closest to the global maxima in the network. (ii) The scatter plot shows the continuity between the input strength-independent subthreshold and suprathreshold resonances in the single PC. (**D**) Response of a minimal PC/IN network with one PC and one IN. Strength of the PC → IN synapse was increased so that a spike in the single PC elicited a spike in the IN, and the background noise was removed for comparison with the resonant response of the single PC. (i) PC firing rate profiles in the minimal network given sinusoidal inputs with suprathreshold input strength *r_inp_* = .3, .8,1 kHz. Feedback inhibition suppressed the resonant peak at 2 Hz and led to the response at higher frequency being resonant (marked with a closed circle). (ii) Scatter plot showing that the FR resonance in the PC (marked with a blue circle) and IN (marked with a red triangle) occur at the same input frequency and scale with the strength of input. (iii) Scatter plot showing that the FR resonant frequency decreases as the duration of inhibition increases. (**E**) Response of the full PC/IN network. Background noise was removed for comparison with the single PC and minimal network cases. (i) PC firing rate profiles in the full network given sinusoidal inputs with suprathreshold input strength *r_inp_* = .3, .8,1 kHz. Feedback inhibition driven by a population of PCs produced lower time-averaged firing rates in the average cell and increased the resonant frequency; the increase in resonant frequency may be due to a mechanism similar to that explored in [18]. (ii) Scatter plot showing that FR resonant frequencies in the PC (marked with a blue circle) and IN (marked with a red triangle) populations increase with input strength. Comparison to the profiles in (i) reveals that the FR resonant frequencies of PC and IN populations separate when the network moves from a band-pass to all-pass regime (i.e., with natural oscillation generated by equal-strength asynchronous drive). (iii) Scatter plot showing that FR resonant frequencies decrease in the full network as the duration of inhibition increases.

**S2 Fig.**
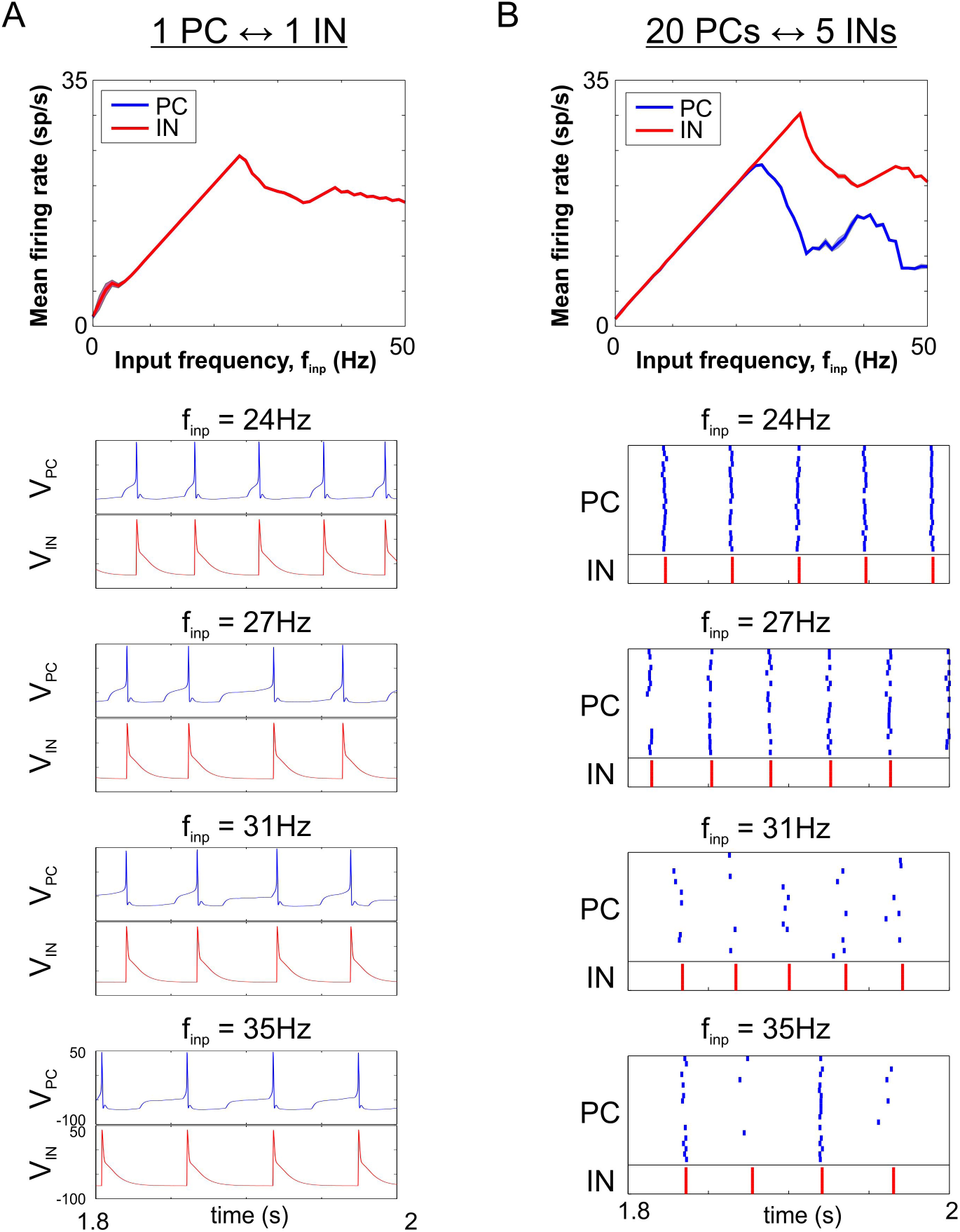
Separation of 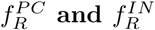requires a PC population with heterogeneous spike times. (**A**)Time-averaged firing rate profiles of PC (blue) and IN (red) cells in a minimal PC/IN network with one PC and one IN given high synchrony, square wave input. The IN can spike only when the PC spikes which causes their firing rates to peak in response to the same input frequency. (**B**) Time-averaged firing rate profiles of PC (blue) and IN (red) populations in the full PC/IN network given high synchrony, square wave input. The IN cells continue spiking after the firing peaks in the PC population because even a subset of PCs spiking on every cycle of the input is sufficient to engage all the INs.

**S3 Fig.**
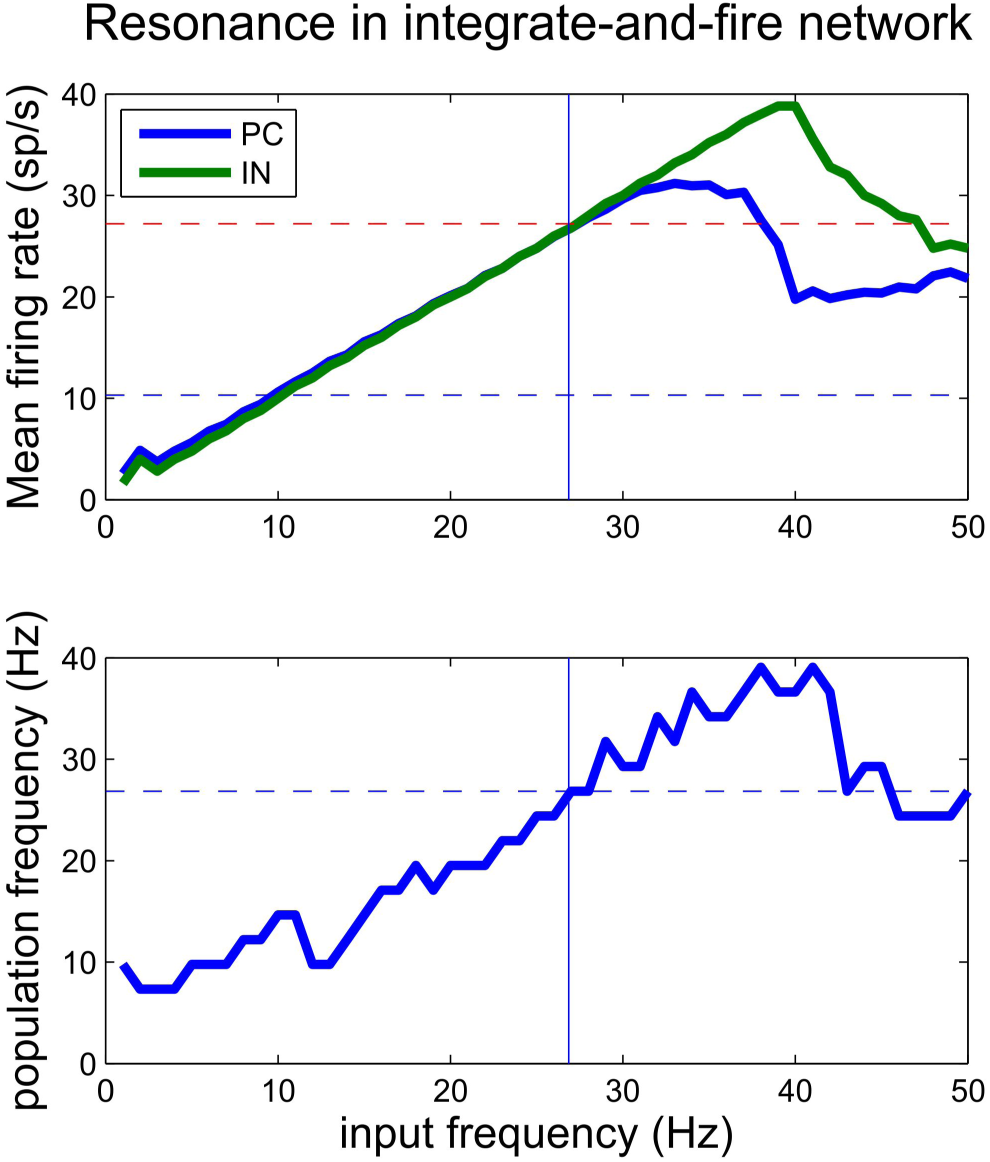
Input frequency-dependent output response profiles in integrate-and-fire networks. Qualitative features of the PFC network model were reproduced in a simpler PC/IN network model with leaky integrate-and-fire (LIF) neurons. (**top**) Firing rate profile for PC (blue) and IN (red) populations. (**bottom**) Population frequency profile for PC and IN populations. Peak population frequency occurs at the input frequency maximizing IN activity (i.e., feedback inhibition). LIF neurons were modeled with membrane potential *V* (mV) governed by: 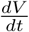 = –*I_inp_*(*t*, *V*) – *gl*(*V* – *E_l_*) – *g_Syn_*(*V* – *E_syn_*) where *t* is time (ms), *g_l_* = 0.1, *E_l_* = –65 mV, *I_inp_*(*t*, *V*) is an excitatory current (μA/cm^2^) reflecting inputs from external sources described in the Methods section, and *I_syn_* denotes synaptic currents (μA/cm^2^) with double exponential conductances driven by other populations. When the membrane potential reaches the threshold of 0 mV, the voltage is reset and held at −65 mV for a refractory period of 3 ms. There were 25 PCs and 5 INs. For PCs, synaptic inputs were inhibitory with *g_syn_* = 0.1, *E_syn_* = –80 mV, 2 ms decay and 0.4 rise time constants. For INs, synaptic inputs were excitatory with *g_syn_* = 0.03, *E_syn_* = 0 mV, 10 ms decay and 0.2 rise time constants. Inputs to the LIF network were the same as the more detailed PFC network described in the Methods section except that *g_inp_* = .00375 mS/cm^2^ and *g_noise_* = 0.0056 mS/cm^2^.

**S4 Fig.**
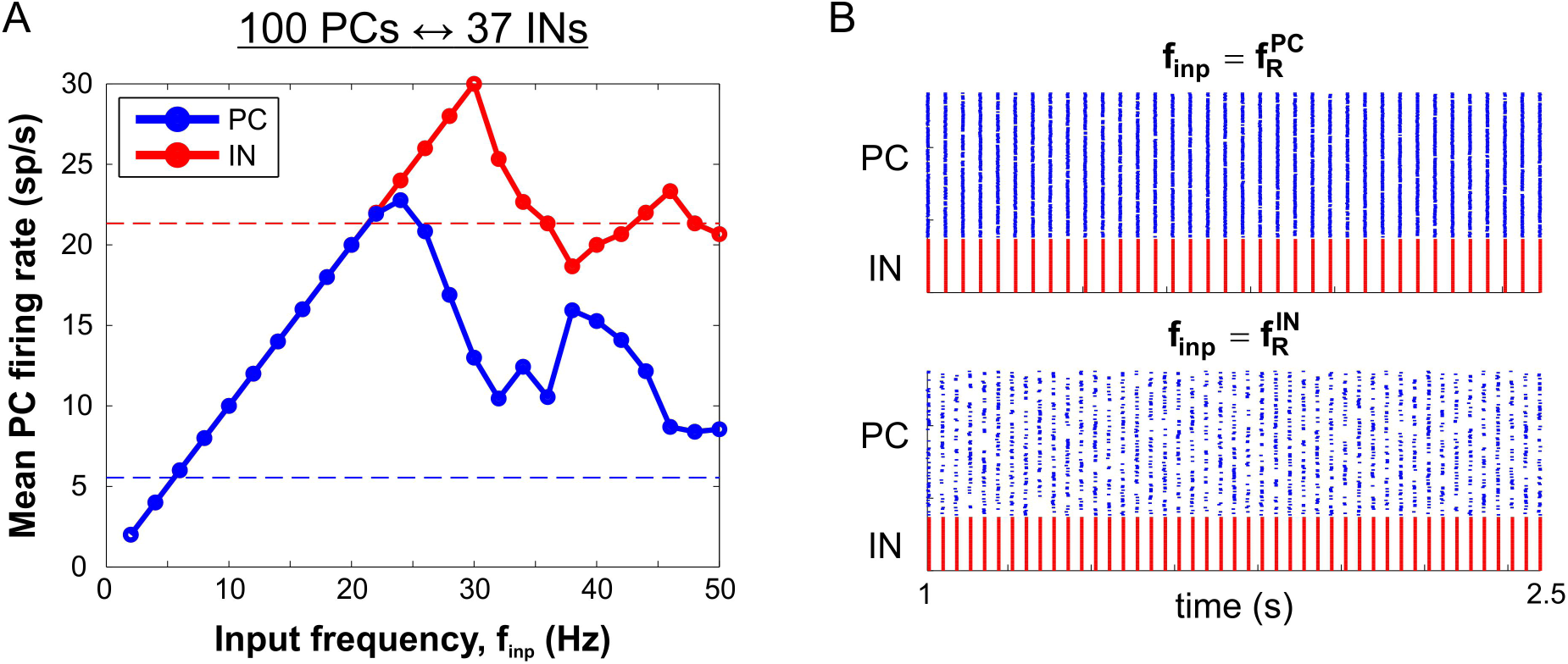
Time-averaged firing rate resonance in the PFC network with 100 PCs and 37 INs. (**A**) Firing rate profile for PC (blue) and IN (red) populations in a PFC network with 100 PCs and 37 INs. All parameters of the model were kept fixed relative to the control model except the number of cells per population was increased. Note that the resonant frequencies are the same as in the control model with 20 PCs and 5 INs. (**B**) Raster plots showing maximal PC spiking at *f_inp_* =
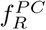and maximal IN spiking (with less PC spiking) at *f_inp_* =
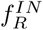.

**S5 Fig.**
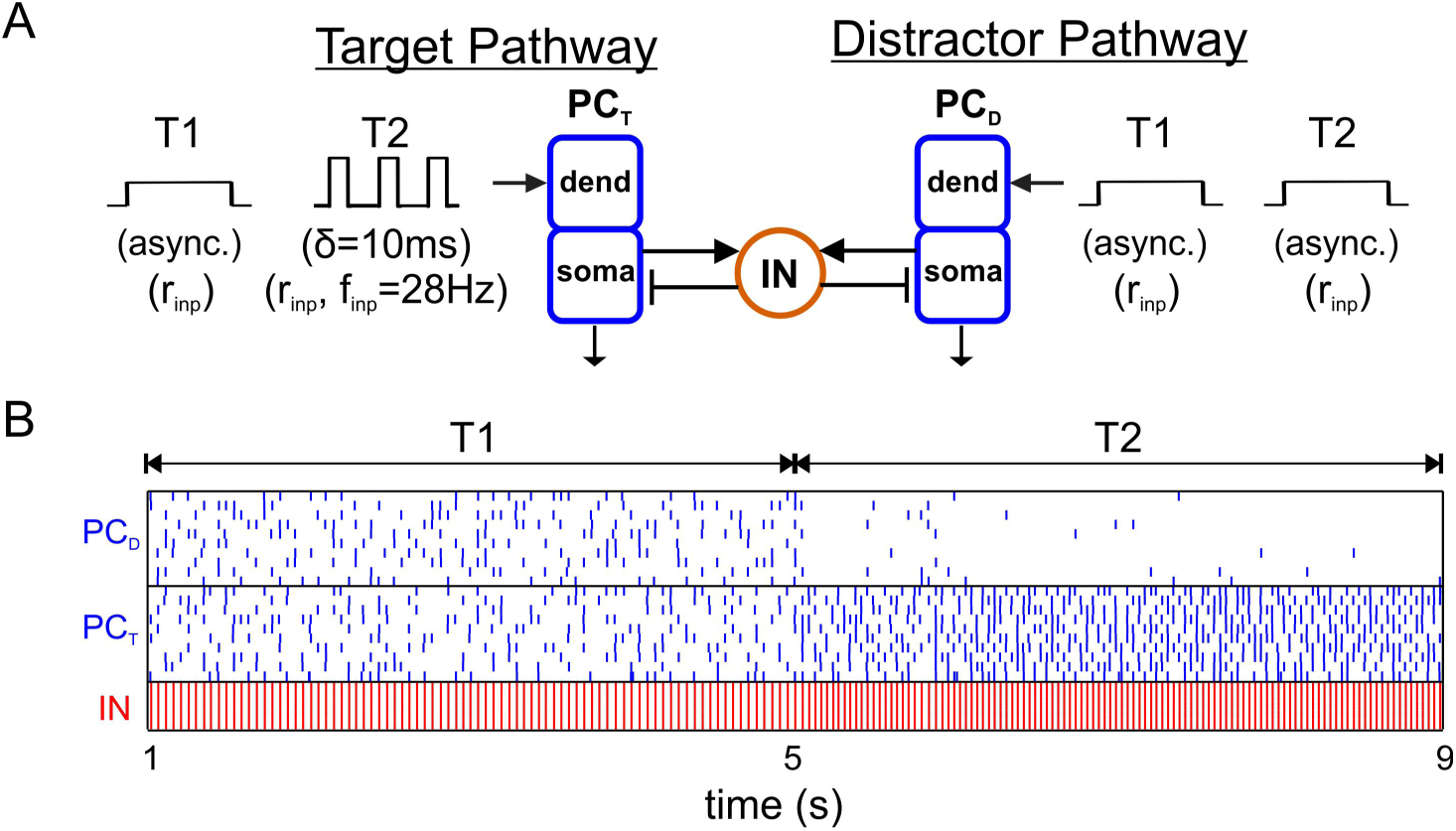
Suppression of response to asynchronous activity occurs within one cycle of the target oscillating more quickly. (**A**) Diagram showing a target PC population, PC_T_, driven by an asynchronous input during period T1 then a medium-synchrony oscillatory input at the 28 Hz *f_pop_*-resonant frequency during period T2 in competition with a distractor PC population, PC_D_, driven by an equal-mean asynchronous input during both periods. (**B**) Raster plot showing that no suppression of either population occurs when their population frequencies are the same during period T1 but that PC_D_ is suppressed within a cycle of PC_T_ oscillating more quickly during period T2.

**S6 Fig.**
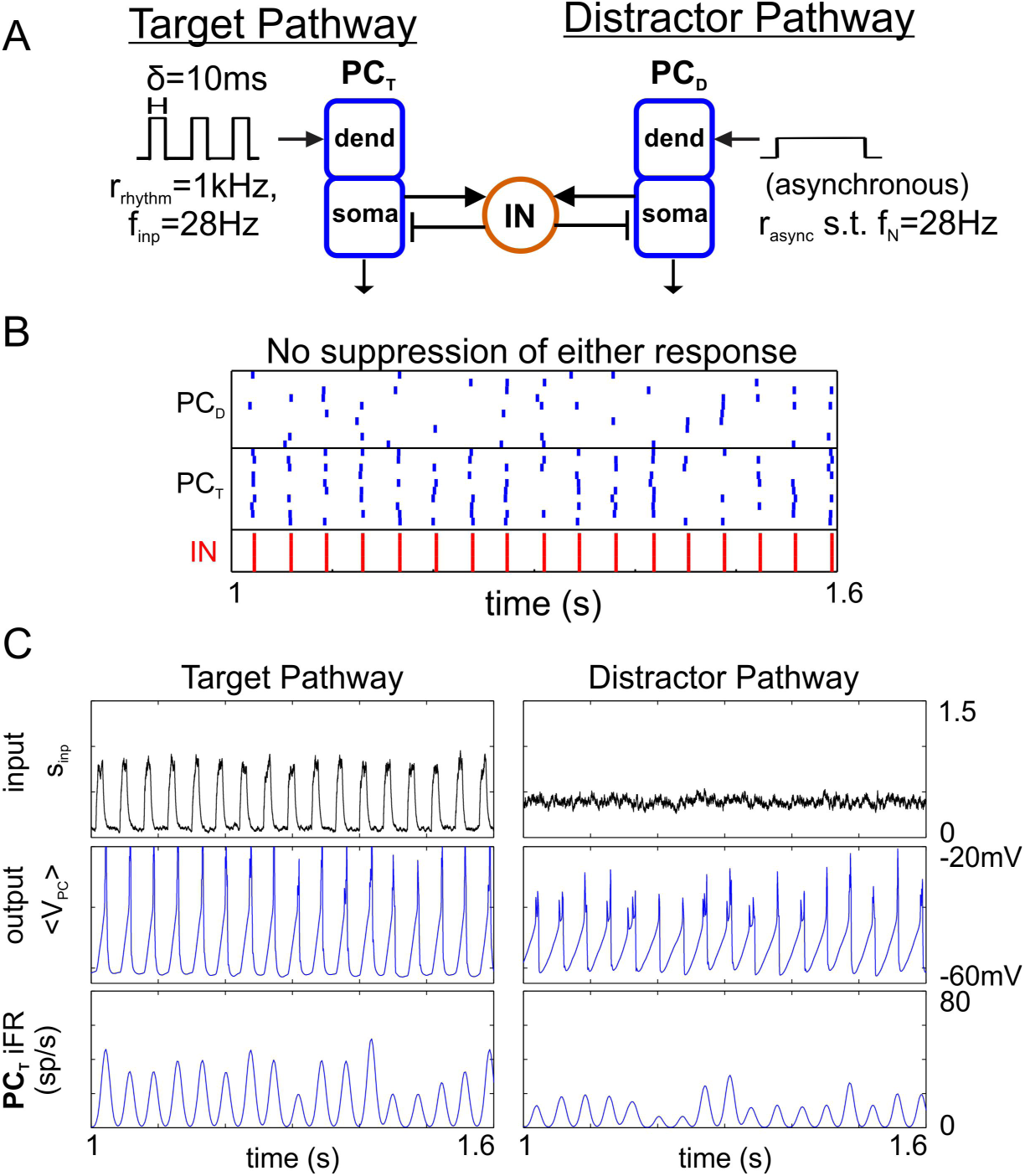
Suppression of response to asynchronous activity does not occur when the natural frequency equals the population frequency of the target. (**A**) Diagram showing a target PC population, PC_T_, driven by medium-synchrony oscillatory input at the 28 Hz *f_pop_*-resonant frequency in competition with a distractor PC population, PC_D_, driven by a higher-rate asynchronous input that produces a 28 Hz natural oscillation. (**B**) Raster plot showing that synchronous spiking occurs in both populations with time-averaged firing rates that would be expected in each population given their inputs in the absence of competition. (**C**) Plots showing the (top) input, (middle) mean population voltage, and (bottom) instantaneous firing rate for (left) PC_T_ and (right) PC_D_ populations.

**S7 Fig.**
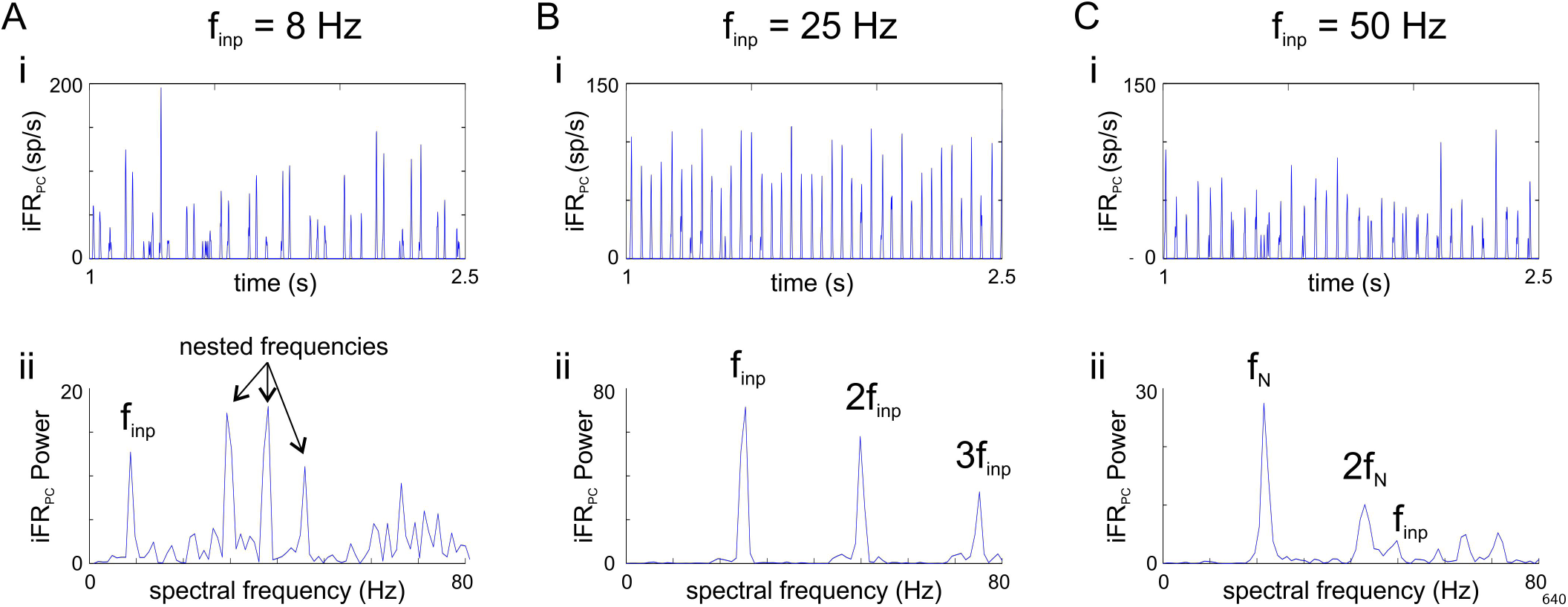
Example iFR power spectra used for the determination of population frequency. (**A**) Response to 8 Hz sinusoidal drive. (i) Nested oscillations in the PC iFR. (ii) Power spectrum with peaks at both the external driving frequency and frequency of internally-generated, nested oscillations. Low-synchrony square wave inputs can also produce nested oscillations. Across realizations, different frequencies may have peak power, which results in ambiguity when *f_pop_* is defined as the frequency with peak power. However, this does not affect the current study because nesting only occurs at frequencies well below the time-averaged firing rate peaks of ongoing inhibition-based oscillations investigated in this work. (**B**) Response to 25 Hz sinusoidal drive. (i) PC population with instantaneous firing rate locked to the period of the input; this occurs for intermediate frequencies of a sinusoidal input and all frequencies of a high-synchrony square wave input up to the IN firing rate resonant frequency. (ii) Power spectrum with peaks at the external driving frequency and its harmonics. (**C**) Response to 50 Hz sinusoidal drive. (i) PC population with instantaneous firing rate paced by the network’s internal time constants; this occurs for driving frequencies above the IN firing rate resonant frequency. (ii) Power spectrum with prominent peaks at the internally-generated, natural frequency and its harmonics as well as a much smaller peak at the external driving frequency. The response to 50 Hz square wave drive, independent of the degree of input synchrony, exhibits a similar asymptotic behavior.

**S8 Fig.**
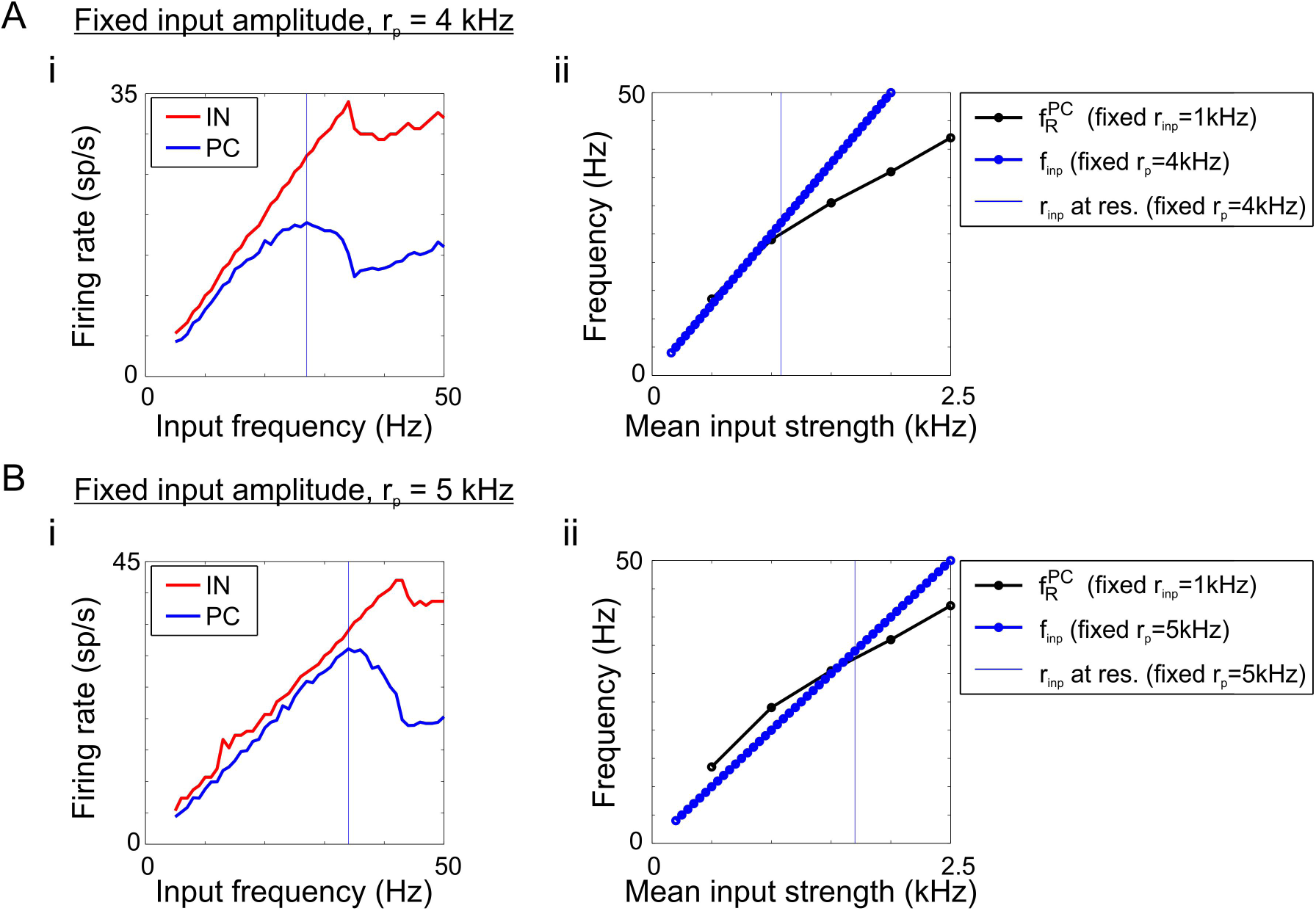
Relationship between resonance with fixed-mean square waves and responses to fixed-amplitude square waves. (**A**) Response to fixed-amplitude square wave with pulse amplitude fixed to *r_p_* = 4 kHz for all driving frequencies. (i) Firing rate (FR) profile for medium-synchrony square waves. Given fixed pulse amplitude, mean input strength increases with input frequency, proportionally to (pulse amplitude) × (inter-pulse frequency) × (pulse width), as an increasing number of 4 kHz pulses occur in the same period of time; 4 kHz corresponds to the amplitude of a fixed-mean square wave pulse when *r_inp_* = 1 kHz and *f_inp_* = 25 Hz. FRs peak at higher frequencies than in response to a fixed-mean square wave. (ii) Plot showing (1) the resonant frequency of peak PC FR given fixed-mean square waves for different drive strengths, *r_inp_* (black), (2) the fixed-amplitude input frequencies corresponding to different drive strengths (blue), and (3) a vertical line marking the fixed-amplitude drive strength at the first peak of PC FR in (Ai). The intersection of these curves shows that the first peak in the fixed-amplitude FR profile in (Ai) occurs when the mean strength for a given *f_inp_* establishes an input strength-dependent FR resonant frequency (determined using fixed-mean square waves) that matches the input frequency. (**B**) Same as (A) except the pulse amplitude was fixed to *r_p_* = 5 kHz for all driving frequencies. 5 kHz corresponds to the amplitude of a fixed-mean square wave pulse when *r_inp_* = 1 kHz and *f_inp_* = 20 Hz. (i) Firing rates peak at an even higher input frequency. (ii) Compared to (Aii), the blue curve shifted to the right because every frequency is associated with a higher mean firing rate when the pulse amplitude increases. The frequency at which the peak occurs in (Bi) corresponds to the mean input strength marked with a vertical line. As in (A), the intersection of the three curves shows that the first peak in the fixed-amplitude FR profile in (Bi) occurs when the input frequency equals the FR resonant frequency given the mean input strength associated with a fixed-amplitude square wave at that input frequency. These results suggest that when pulse amplitude is fixed, as in the fixed-mean case, the resonant frequency increases with mean input strength. However, the mean input strength increases with input frequency when pulse amplitude is fixed, leading to a more complicated relationship between input frequency and local maxima in the response profile for fixed-amplitude square wave inputs.

## Acknowledgments

We would like to acknowledge Horacio Rotstein for his feedback and generous conversations about resonance as well as Michelle McCarthy and Dan Bullock for their insights during our discussions of the results.

